# Loss of E3 ligase *HvST1* function substantially increases recombination

**DOI:** 10.1101/2023.05.19.541444

**Authors:** Jamie Orr, Sybille Mittmann, Luke Ramsay, Dominika Lewandowska, Abdellah Barakate, Malcolm Macaulay, Nicola McCallum, Robbie Waugh, Isabelle Colas

**Author notes:** **Corresponding author:** Isabelle Colas **Email:**. ***Authors contributed equally to this work*** (Jamie Orr, Sybille Mittmann).

## Abstract

During meiosis, genetic recombination occurs via repair of DNA double-strand breaks (DSBs) as crossovers (COs) resulting in the exchange of parental genetic material (De Muyt *et al.*, 2009). Crossovers are important for chromosome segregation and shuffling genetic variation, but their number and distribution are tightly regulated (Zickler and Kleckner, 2015). In barley and other large genome cereals, recombination events are limited in number and mainly restricted to the ends of chromosomes (Mascher *et al.*, 2017), constraining progress in plant breeding. Recent studies have highlighted subtle differences in meiotic progression (Higgins *et al.*, 2012; Phillips *et al.*, 2013) and the distribution of recombination events in barley compared to other plants (Colas *et al.*, 2016; Colas *et al.*, 2017; Colas et al 2019), indicating possible evolutionary divergence of the meiotic program in large genome crops. Here we identify a spontaneous loss of function mutation in the grass specific E3 ubiquitin ligase *HvST1* (*Sticky Telomeres 1*) which results in semi-sterility in barley. We show that abnormal synapsis in the absence of HvST1 function increases overall recombination by up to 2.5-fold and that HvST1 is capable of ubiquitinating ASY1, a key component of the lateral elements of the synaptonemal complex. Our findings shed light on an evolutionarily divergent pathway regulating synapsis and recombination in cereals. This natural loss of function variant presents new opportunities for the modulation of recombination in large genome cereals.

## Introduction

Homologous recombination (HR) through resolution of double strand breaks (DSBs) as crossovers (COs) is an important trait for breeders, who rely on this mechanism of genetic exchange during meiosis to create novel crop varieties. In barley, HR occurs mainly in distal chromosome regions leaving large sections of the genome almost devoid of recombination (Kunzel *et al.*, 2002; Lambing *et al.*, 2018). In plants, pairing of replicated homologous chromosomes in meiosis I is guided by the formation of DSBs catalyzed by SPO11-1 and SPO11-2 (Grelon *et al.*, 2001; Hartung *et al.*, 2007). One strand of DNA at each side of the DSB is partially degraded (resected) by the MRE11-Rad50-NSB1 complex leaving 3’ single stranded DNA (Daoudal-Cotterell *et al.*, 2002; Wang *et al.*, 2014). The RECA protein family, which includes RAD51 and DMC1 proteins, mediate strand invasion to create a “D-loop” (Pradillo *et al.*, 2012; Da Ines *et al.*, 2013). It is postulated that D-Loops in plants are resolved through double-Holliday junction (<u>dHj</u>) recombination intermediates to make COs (Neale *et al.*, 2006) via one of two pathways, resulting in either class I or class II COs. Class I CO resolution depends on the ZMM proteins (Zip1-4, Msh4/Msh5, and Mer3) (Higgins *et al.*, 2005; Higgins *et al.*, 2008b) and is subject to CO interference, a phenomenon that restricts the proximity of DSBs repaired as COs via this pathway (Capilla-Perez *et al.*, 2021). Class II CO resolution is thought to depend on the MUS81 pathway (Higgins *et al.*, 2008a) and is not subject to CO interference (Falque *et al.*, 2009). In Arabidopsis and barley, it is estimated that only 5% of DSBs are resolved as COs (Serra *et al.*, 2018, Higgins *et al.*, 2012) and that 85% of these are resolved through the interference sensitive class I pathway. This introduces significant linkage drag which limits the ability of barley breeders to separate desirable and undesirable traits. The barley secondary gene pool—land races and wild relatives—are a valuable resource to breeders for novel traits such as disease or drought resistance. However, linkage drag can impose a severe yield or quality penalty when attempting to transfer these traits to elite varieties. Therefore, the ability to increase recombination and break interference is appealing to plant breeders.

At the onset of meiosis in barley the telomeres cluster to one side of the nucleus and the formation of the synaptonemal complex (SC, synapsis) is initiated (Higgins *et al.*, 2012, Colas *et al.*, 2017). The SC is a tri-partite structure consisting of two lateral elements which form along replicated sister chromatids, helping to constrain them, and a central element which, beginning from the clustered telomeric region, forms between the lateral elements helping to physically link the paired homologous chromosomes during prophase I (Orr *et al.*, 2021a). The SC is later dissolved prior to the first meiotic division, leaving the chromosomes held together by chiasma—the cytogenetic manifestation of COs—which help to ensure correct chromosome segregation during the second round of meiotic division (Orr *et al.*, 2021a). Synapsis can be visualized using antibodies against key SC proteins such as ASY1, a component of the lateral elements, and ZYP1, a component of the central element (Colas *et al.*, 2017). Impairing the recombination pathway not only affects crossover outcomes but can also affect synapsis, demonstrating the tight interplay of the two parallel mechanisms (Grey *et al.*, 2022). However, similar disruptions to these pathways can result in substantial differences in meiotic phenotype in different plant species. For example, barley desynaptic *des10*, which carries a mutation in MLH3 (Colas *et al.*, 2016) displays abnormal synapsis contrary to the equivalent mutation in Arabidopsis (Jackson *et al.*, 2006; Colas *et al.*, 2016).

Recent studies have reported that mutations affecting the function or expression of *FANCM*, *RECQ4* and *HEI10* genes have the potential to increase recombination (Serra *et al.*, 2018; Mieulet *et al.*, 2018; Arrieta *et al.*, 2021) which is of great interest in large genome crops such as barley where the distal bias for recombination events is particularly pronounced (Kunzel *et al.*, 2002; Lambing *et al.*, 2018). To this end, we have exploited a collection of 15 *desynaptic* mutants that have been initially cytologically classified by their chromosome behaviour at metaphase I in the early 1970s (Hernandes-Soriano, 1973). Here we show that one of these lines—BW233—is mutated in the RING domain of a grass specific E3 ubiquitin ligase, that we call *HvST1* (*Sticky Telomeres 1*). We have shown that HvST1 is a functional E3 ubiquitin ligase and is capable of ubiquitinating ASY1 in vitro. Despite an abnormal synapsis progression, we found that loss of E3 ubiquitin ligase activity in *Hvst1* leads to an overall increase recombination which could be exploited during breeding programs.

## Materials and Methods

### Plant material

*H. vulgare* BW233 is one of a collection near isogenic lines (NILs) derived from a collection of 15 *desynaptic* mutants that have been initially cytologically classified by their chromosome behaviour at metaphase I in the early 1970s (Hernandes-Soriano, 1973). These mutants have been backcrossed to a common barley cultivar Bowman to create NILs and initially genotyped to find the causal mutation for their meiotic phenotype (Lundqvist *et al.*, 1997; Druka *et al.*, 2010). Barley cv. Bowman and NIL BW233 plants were grown under the following conditions: 18-20°C for 16 hours of light and 14-16°C for 8 hours of dark. For cytology, spikes of 1.4 to 2.5 cm were collected, and anthers were prepared as previously described (Colas *et al.*, 2016) For fine mapping, 12 seeds from each of the 16 families of the F_2_ population of BW233 x Barke were grown in 24-pot trays. For the F_3_ recombination assay, 48 families of the harvested F_2_ BW233 x Barke population plants were chosen according to their genotypes across the *des12* interval. For the cultivar Barke and BW233 genotypes, 8 seeds per family were sown and grown as described for the F_2_ population (24 families per genotype). DNA was extracted from young leaf tissue (two-leaf stage) using the Qiagen DNeasy 96 well plate Kit with the automated station QIAxtractor® or QIAcube®. RNA was extracted from 50 anthers in prophase I, using the Qiagen RNeasy Mini Kit. RNA quantity and quality were assessed with the NanoDrop 2000.

### Cytology

Metaphase spreads were performed as described by Higgins *et al*. (2012) with minor changes. Briefly, fixed anthers were incubated in 1% cellulase and 2% Pectolyase solution at 37°C for 30 minutes. The reaction was stopped by replacing the enzyme mix with cold (4°C) sterile distilled water. Nuclei preparation for immuno-cytology and Immuno-FISH was performed as previously described (Colas *et al.*, 2016)

#### Fluorescence in situ hybridization (FISH)

Probes for Centromeres (Bac7) and Telomeres (HvT01) were amplified by PCR and labelled by nick translation with Alexa-Fluor as previously described (Colas *et al.*, 2016). Hybridization mix and denaturation was performed as previously described (Arrieta *et al.*, 2021). Slides were washed in 2xSSC at room temperature (RT) for 10 minutes before digestion with 0.01% (w/v) pepsin in mildly acidified SDW for 45 to 90 seconds at 37°C. Slides were dehydrated using a series of ethanol concentrations (50%, 70%, 90% to 100%) for 2 minutes each and left to air-dry at RT. Hybridization mix (40 µl) was applied to the slides, and slides were incubated at 75°C for 4 minutes, then over night at 37°C in a dark humid chamber. Slides were washed and prepared as previously described (Colas *et al.*, 2016)

#### Immunocytology

Slides were prepared as previously described (Colas *et al.*, 2016). The primary antibody solution consisted of one or multiple antibodies; anti-AtASY1 (rabbit, 1:1000), anti-AtZYP1 (rat, 1:500), anti-AtDMC1 (rabbit, 1:500), anti-HvMLH3 (rabbit, 1:500), anti-HvHei10 (rabbit, 1:500), diluted in blocking solution (1X PBS with 5% Donkey/Goat serum). For Immuno-FISH, slides were washed in 2x SSC for 5 minutes after the secondary antibody incubation, followed by 10 minutes in 1x PBS. Slides were then fixed with 1% PFA for 10 minutes, rinsed briefly in 1xPBS before following the FISH protocol described earlier from the dehydration step.

#### Microscopy and Imaging

3-dimensional Structured Illumination Microscopy (3D-SIM) and confocal images were acquired on a DeltaVision OMX Blaze (GE Healthcare) and LSM-Zeiss 710 respectively, as previously described (Colas *et al.*, 2016). Images were de-convolved with Imaris deconvolution module Clearview 9.5 and processed for light brightness/contrast adjustment with Imaris 9.5 (Randall *et al.*, 2022).

### Positional gene identification

Initial genetic mapping used a custom 384 SNP genotyping array using the Illumina beadXpress platform on an F_2_ segregating population derived from a cross between BW233 (*des12.w*) and *cv*. Morex, using the segregation of the semi-sterile phenotype of *des12.w* as a Mendelian trait (Druka *et al.*, 2010). The *des12.w* interval was delineated a 1.0 cM interval on chromosome 7HL, by SNP markers found in MLOC_38602 and MLOC_67621. The region was further delineated using custom designed KASPar SNP assays on 960 plants of the segregating F_2_ population. KASPar assays were designed based on SNPs extracted from the manifest files of a 9k iSelect Genotyping platform developed at the James Hutton Institute (Bayer *et al.*, 2017). Synteny with rice and *Brachypodium* was used to identify genes that would be useful for delineation of the interval and contigs from cvs. Morex, Bowman and Barke. Assemblies were pulled out using the cDNA of the gene sequence, gathered from Ensembl Plants (Bolser *et al.*, 2017) and BARLEX (Colmsee *et al.*, 2015). BACs that contained the 9 significant genes in the *des12.w* interval were obtained from the HarvEST web server (Close *et al.*, 2009) to find sequences of interest to BLAST, sequence, and identify any polymorphisms between BW233 and WT alleles (from cultivars Bowman, Barke, Morex, Freya, and Glages). Linkage maps were constructed using the combined genotypic and phenotypic data described above using JoinMap 4.1 (Stam, 1993).

### F_3_ recombination assay

The F_3_ recombination assay was based on KASP™ genotyping chemistry using 48 markers spanning 3 chromosomes ( 1H, 5H, and 6H; Table S 1 ). The selected markers and frozen leaf punches were sent to LGC for KASP assay development. Data was visualised with Flapjack (Milne *et al.*, 2010) and further analysed in Microsoft Windows Excel 2013. Recombination frequencies were calculated to generate linkage maps with MapChart (Voorrips *et al.*, 2002). A subset of the F_3_ families was s u b s e q u e n t l y analysed with the barley Illumina iSelect 50k SNP array (Bayer *et al.*, 2017). Monomorphic markers were removed, and a sliding window was used to correct for poorly mapped markers. F_2_ recombination events, visible as large monomorphic blocks, generated unusually high crossover counts for individual markers, markers with crossover counts above 5 across all 95 plants were removed to account for this. All code used in 50K analysis and plotting is available online (see code availability statement).

### Gene model and expression

The *HvST1* gene model was constructed using the Barley Reference Transcriptome (BaRT) v2.18 (Coulter *et al.*, 2022). HvST1 meiotic expression was plotted using data from the Barley Anther and Meiocyte Transcriptome (BAnTr) (Barakate *et al.*, 2021).

### Gene Cloning

*HvST1,* truncated *Hvst1*, and *HvASY1* cDNA was synthesised with the SuperScript® III First-Strand Synthesis System (Invitrogen) using the included oligo(dT) primer. cDNA was then amplified using nested PCR (Primers in Table S2) with High-Fidelity Phusion or Q5 DNA polymerase (New England Biolabs), and the PCR products ligated into pGEM®-T Easy vector (Promega). Competent cells of *E*. *coli* JM 109 were transformed with the ligations and plated on solid LB medium containing Ampicillin, X-gal, and IPTG for selection and blue-white screening. Plasmid DNA was purified from positive clones using the QIAprep Spin Miniprep Kit (Qiagen). The cDNA sequence was verified by Sanger sequencing using T7 and SP6 primers.

### Protein Expression Constructs

*HvST1,* truncated *Hvst1*, and *HvASY1* coding regions were amplified from respective pGEM-T plasmids by PCR using primers with attB1 and attB2 Gateway sites (Table S3). The TEV protease cleavage site was introduced by overlapping TEV-oligonucleotide and the PCR product cloned into pDONR207 using BP clonase (Invitrogen). The resulting BP clonase reaction was transformed into *E*. *coli* DH5α chemical competent cells (ThermoFisher Scientific). The transformation was plated on LB agar plates containing 25 µg/ml gentamycin. After colony PCR screening with attB1 and attB2 primers, overnight cultures of the positive clones were prepared in LB medium with 25 µg/ml gentamycin and their plasmid DNA prepared and verified by Sanger Sequencing using PDONR207 forward and reverse primers. The *HvST1* insert was then transferred into pDEST-HisMBP (Nallamsetty *et al.*, 2009) using LR clonase (Invitrogen)^21^. The LR clonase reaction was used to transform *E*. *coli* DH5α competent cells before plating on LB agar plates supplemented with 100 µg/ml ampicillin. Plasmid DNA of positive clones was prepared and checked by restriction digest using HincII for *HvST1/Hvst1* and BamHI and BglII (New England Biolabs) for *HvASY1*. The selected clones were then transferred into *E*. *coli* Rosetta™ 2(DE3) pLysS (Novagen). The transformations were plated on LB agar plates supplemented with 100 µg/ml ampicillin and 34 µg/ml chloramphenicol. A single colony was grown overnight in LB medium with both antibiotics.

### Protein Expression and Purification

Single colonies from successful transformants were grown to OD600 nm of approximately 0.6 in LB with of 100 µg/ml Ampicillin and 34 µg/ml Chloramphenicol at 37°C and 220 rpm. Cultures were cooled on ice before induction with 0.1 mM IPTG and grown overnight at 18°C and 220 rpm. Induced cells were then pelleted by centrifugation, flash frozen in liquid nitrogen, and stored at -80°C until use. Induced cell pellets were lysed in BugBuster® Master Mix (Novagen) with cOmplete™ the EDTA-free Protease Inhibitor Cocktail (Roche). The overexpressed protein was purified using HisPur NiNTA resin (ThermoFisher). For removal of the HisMBP tag, NiNTA elution buffer was exchanged with 50 mM TrisHCl, pH 8.0 using Pierce 10K MWCO spin columns (ThermoFisher) before digestion with ProTEV Plus (Promega) overnight at 4°C with the addition of 1% Triton X-100. Detergent was removed using a Pierce detergent removal spin column (ThermoFisher), then a second round of NiNTA capture removed the cleaved tag, leaving the purified protein in the spin column flow through. Purified protein was exchanged into PBS (pH 7.4) using Slide-A-Lyzer G2 dialysis cassettes (ThermoFisher) and stored at 4°C until use. The concentration of the purified protein was measured using the Pierce 660 nm microplate assay (ThermoFisher) and absorbance read on the Varioskan Lux (ThermoFisher).

### *In vitro* ubiquitination assays

Purified HvST1 interactions with E2 conjugating enzymes were determined by screening against a panel of human E2 conjugating enzymes using the E2-scan plate (Ubiquigent). Reactions were evaluated for accumulation of polyubiquitinated products via SDS-PAGE followed by staining with SimplyBlue Safe Stain (ThermoFisher) and western blotting with mouse anti-ubiquitin conjugate antibody at a dilution of 1:10000 (Ubiquigent). Subsequent ubiquitination time course assays were conducted using histidine tagged UBE1, UBE2D4, bovine ubiquitin, and 8 mM ATP (Ubiquigent). HvST1 autoubiquitination time course assays and substrate (HvASY1) ubiquitination assays were incubated at 37°C for between 0 and 60 minutes. All control reactions were incubated at 37°C for 60 minutes.

These reactions were analysed via SDS-PAGE followed by SimplyBlue Safe Stain (ThermoFisher) and western blotting using mouse anti-ubiquitin conjugate antibody as above, rat anti-HvST1 antibody (1:200), mouse anti-^6^His antibody (1:10000; ThermoFisher), and rabbit anti-HvASY1 antibody (1:5000; Agrisera; this study; AS21 4690). Excised gel fragments from autoubiquitination experiments were prepared as previously described (Lewandowska *et al.*, 2019). Mass spectrometry was carried out by Dundee University FingerPrints Proteomics Facility. The resulting raw files were processed and searched using MaxQuant (v1.6.10.43) (Tyanova *et al.*, 2016) and the Andromeda peptide search engine (Cox *et al.*, 2008, Cox *et al.*, 2011) against the E. coli BL21 RefSeq proteome (GCF_014263375.1) combined with a custom set of sequences for the purified proteins added to the assay.

### TUBE capture

GST-tagged tandem ubiquitin binding entities (GST-TUBEs) were expressed and purified as described by Skelly *et al*. (2019) The HvASY1 ubiquitination assay was prepared as above and incubated for two hours at 37°C. The reaction was passed through a 50 KDa size exclusion column (ThermoFisher) to eliminate remaining free ubiquitin. GST-TUBE was added to the column retentate to a final concentration of 200 µg/ml before overnight glutathione agarose (ThermoFisher) capture at 4°C with gentle agitation. Each fraction of the glutathione agarose capture was retained and of equal volume. The captured protein was divided in two and half was incubated for two hours at 37°C with broad spectrum deubiquitinating enzyme (DUB; LifeSensors). The results of the assay were determined via SDS-PAGE and western blotting as described above.

### CRISPR/Cas9 construct

To find suitable CRISPR/Cas9 knockout targets in the *HvST1* gene, the coding sequence was searched using the online algorithm at http://www.broadinstitute.org/rnai/public/analysis-tools/sgrna-design. Two spacer sequences of 20 nucleotide with the highest score and a guanine at their 5’ end as transcription start and preferably an A or G at the 3’ end for increased CRISPR/Cas9 efficiency were selected. A search for potential off-targets was performed using the barley genes database developed at the James Hutton Institute (https://ics.hutton.ac.uk/barleyGenes/index.html). Two complementary oligonucleotides of 20 nucleotides were designed for each target with an additional 4 nucleotides at their 5’-end for cloning into AarI restriction sites in the destination vector pBract214m-HvCas9-HSPT. This vector contains a barley codon optimised Cas9 (HvCas9) under the control of maize ubiquitin promoter and *Arabidopsis* heat shock protein 18.2 terminator and sgRNA scaffold under the control of rice small nuclear RNA (snRNA) U6 promoter (OsU6p). The complementary oligonucleotides (Table S 4 ) were annealed using equal amounts of each oligo. Oligonucleotide mixtures were heated in boiling water then left to cool to room temperature. 1 µl of duplex DNA was ligated into 25 ng of pBract214m-HvCas9-HSPT vector linearized with AarI using the ligation reagents and protocol described in pGEM®-T Easy Vector System kit (Promega). The ligation reaction was incubated overnight at 4°C then transformed into competent *E. coli* DH5α strain (New England Biolabs) and plated on LB medium containing 25 µg/ml Kanamycin. Inserts were verified by colony PCR and Sanger sequencing as described above. Additionally, the integrity of the plasmids was checked by digestion with restriction enzyme BglI (Thermo Fisher Scientific). The plasmid DNA was then transformed into *Agrobacterium tumefasciens* AGL1 strain containing replication helper pSoup (http://www.bract.org/constructs.htm#barley) by electroporation. Briefly, 50 µl electro-competent Agrobacterium AGL1/pSoup was thawed on ice, 1 µl plasmid DNA was added, and the mixture was transferred to an electroporation cuvette and pulsed at 2.5 Kv. 1 ml YEB medium was added and the mixture was transferred into 15 ml falcon tube (Sigma Aldrich) and incubated at 28°C shaking at 200 rpm for 2–3 h before plating on LB agar supplemented with 50 µg/ml Kanamycin (Merck Millipore). The plates were incubated for 2– 3 days at 28°C. Transformed *Agrobacterium* clones from each CRISPR construct were grown in YEB medium and used to transform barley Golden Promise immature embryos (Bartlett *et al.*, 2008) in the plant transformation facility at The James Hutton Institute. Transgenic plants containing CRISPR constructs were regenerated under hygromycin selection.

### CRISPR/Cas9 screening and genotyping

For screening the T0 CRISPR plants, 2mm leaf discs were collected in 20 µl Phire Plant PCR Buffer (Thermo Fisher Scientific) and crushed with a 100 µl pipette tip. For construct CRISPR01 genotyping, the KAPA3G Plant PCR Kit (Sigma-Aldrich) was used to amplify an 824 bp fragment spanning the gene target site using an initial touchdown for 6 cycles with annealing at 70°C decreasing by 1.6°C per cycle followed by 34 cycles annealing at 62.1°C. An amplicon for construct CRISPR02 was amplified with the Phire Plant Direct PCR kit using an initial touchdown annealing for 8 cycles starting at 72°C and decreasing by 0.9°C per cycle followed by 31 cycles annealing at 65.4°C. Products were sequenced by Sanger Sequencing as described above to verify potential mutations. Based on sequencing data, 8 individuals were identified that had mutations at the expected gene location for the CRISPR01 construct. 20 seeds of each were sown in 24-pot trays soil in the glasshouse. At the two-leaf stage, 2 mm leaf discs were collected, and plants were genotyped to identify plants that were homozygous for the identified mutations as described above. Additionally, plants were screened by PCR and gel electrophoresis for the presence or absence of the Cas9 transgene with Cas9-F2 and Cas9-R2 primers (Table S5). Five plants originating from three different T0 plants were identified to be homozygous and contain no Cas9 transgene. These lines were re-potted in 9-inch pots of soil to generate more seeds and tillers for cytology analysis. As control two plants that were found to contain no mutations were also included. Plants were grown for 6 weeks until they reached meiosis and inflorescences were collected for chromosome spreads and immunolocalization as described above.

### Phylogenetic analysis

Proteomes from barley (*H*. *vulgare*), rice (*O*. *sativa*), wheat (*T*. *aestivum*), maize (*Z*. *mays*), tomato (*S*. *lycopersicum*), cassava (*M*. *escualenta*), pineapple (*A*. *comosus*), soybean (*G*. *max*), clementine (*C*. *clementina*), *Brassica oleracea*, Tabaco (*N*. *tabacum*), *Arabidopsis thaliana*, *Amborella trichopoda*, *Physcomitrella patens*, yeast (*S. cerevisiae*), *C*. *elegans*, *D*. *melanogaster*, zebrafish (*D*. *rerio*), human (*H*. *sapiens*), mouse (*M*. *musculus*), and clawed frog (*X*. *laevis*) were clustered into ortholog groups using Orthofinder (Emms and Kelly, 2019; v2.3.3). HvST1 clustered only with proteins from included Poaceae proteomes. Further potential orthologs within Poaceae and the best possible alignments within *Arabidopsis* and monocot spp. were determined by BLAST alignment to the non-redundant archive (Altschul *et al.*, 1990) and larger EggNOG (Huerta-Cepas *et al.*, 2019_;_ v5.0) pre-computed orthologous groups. The longest isoforms of these proteins were aligned using MAFFT (Katoh *et al.*, 2002; v7.266), and maximum likelihood phylogeny constructed using IQ-TREE (Minh *et al.*, 2020; v2.0.3) using model JTT+G4 with ultrafast bootstrapping (n=1000). The resultant phylogeny was visualised using ggtree (Yu, 2020) (v.2.5.1).

## Results

### The *des12.w* phenotype is due to a frame shift mutation in a novel E3 ubiquitin ligase

BW233 *(des12.w)* is semi-sterile isogenic line of the barley cv. Bowman (Fig. 1a) that carries a spontaneous mutation which originated in *cv*. Freja. In contrast to Bowman (wild type; WT) metaphase I (Fig. 1b), BW233 metaphase I (Fig. 1c) chromosomes are sticky and often interlocked in telomeric regions (arrows) and a statistically significant increase in mean rod-bivalent and decrease in mean ring bivalent chromosomes (T-test, Benjamini and Hochberg corrected p = 0.024 for both) are observed (Fig. 1d and Fig. S1). Chiasma counts from Bowman (n=19) and BW233 (n=44) meiocytes at metaphase I showed a significant decrease in BW233 chiasma (Wilcoxon rank sum test, p=0.0055) and altered distribution compared to Bowman (Fig. S2). Chromosomes interlocks often persist at anaphase I (Fig. 1e) and anaphase II in BW233 meiocytes (Fig. 1f), potentially causing lagging chromosomes and mis-segregation.

**Figure 1:**
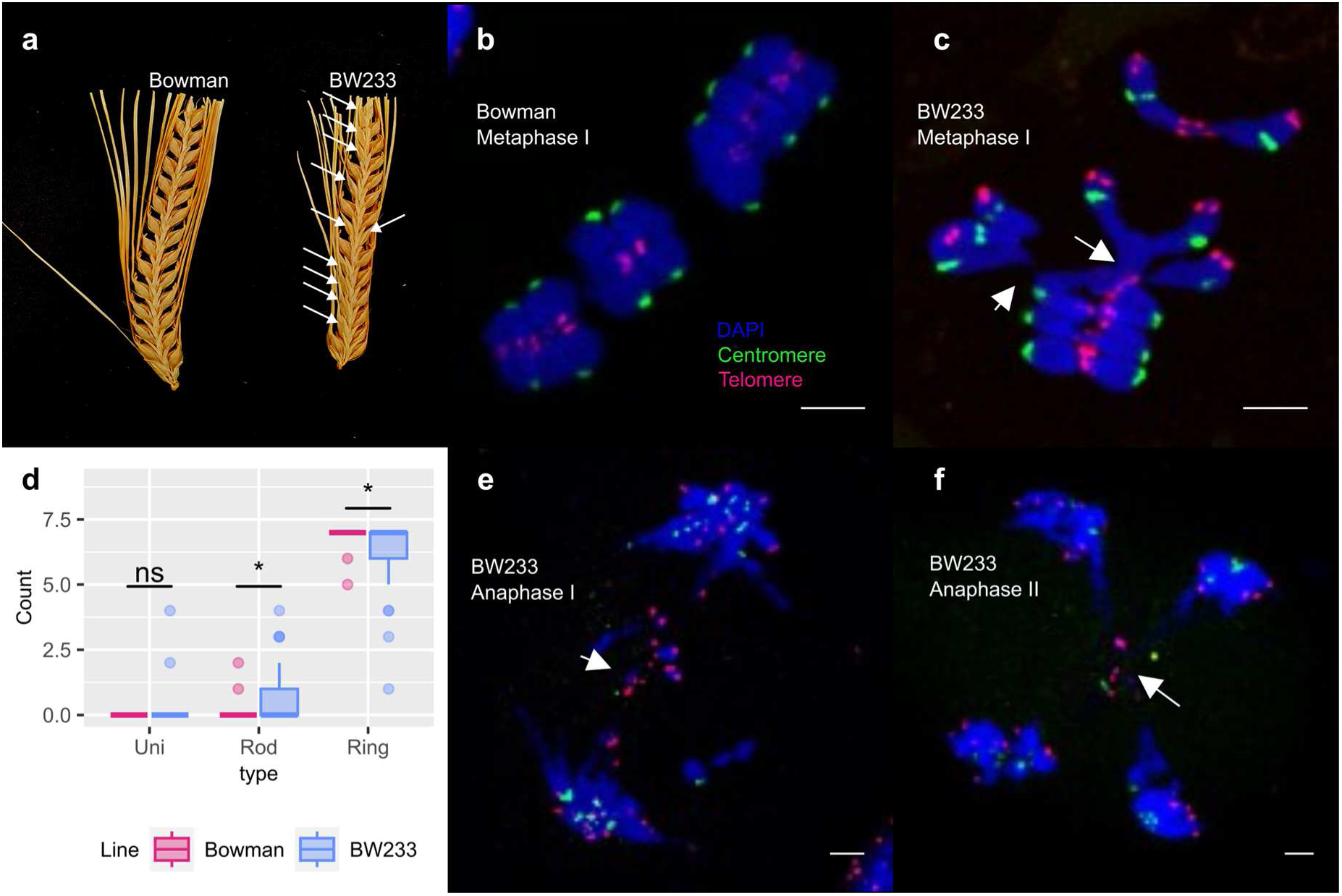
BW233 phenotype. a) Spike fertility comparison between Bowman and BW233, with missing seed indicated by white arrows. b) Bowman metaphase I with 7 ring bivalents. c) BW233 metaphase I, with 7 bivalents often interlocked at the telomere region (white arrows). d) Box plot of univalent, rod bivalent, and ring bivalent counts in *HvST1* and *Hvst1* plants with significance (T-test) indicated above. e) BW233 anaphase I, showing lagging chromosomes and chromosomes bridges (white arrow). f) BW233 anaphase II with interlocked chromosomes at the telomere region (white arrow). DAPI (blue), telomeres (red), centromeres (green). Scale bar 5 µm

To find the causal mutation of the *des12.w* phenotype, we used F_2_ populations of two crosses — BW233 x *cv*. Barke and BW233 x *cv*. Morex — scoring semi fertility as the segregating phenotype (Fig. 1a). We initially mapped the *des12.w* mutation to the long arm of chromosome 7H (Druka *et al.*, 2011) and by positional cloning (Fig. S3) identified a short exonic mononucleotide microsatellite in HORVU7Hr1g092570 (Morex v3: HORVU.MOREX.r3.7HG0722700) (Mascher *et al.*, 2021) that contains an extra guanine in BW233 compared to WT cultivars Bowman, Barke, Freja, Morex, and Betzes, which induces a frameshift (Fig. 2a and Fig. S4). HORVU.MOREX.r3.7HG0722700 is a 990 bp single exon gene encoding a 330 amino acid RING finger domain containing putative E3 ubiquitin ligase (34) that we have named *STICKY TELOMERES 1* (*HvST1*) (Fig. S4). The frame shift in *des12.w* (*Hvst1*) introduces a premature termination codon in *HvST1* that eliminates the RING domain and compromises gene function (Fig. 2a and Fig. S4). In our ‘barley anther and meiocyte transcriptome’ (Barakate *et al.*, 2021) dataset, we found that *HvST1* expression was significantly enriched in staged meiocytes when compared to anthers, rising to peak expression in pachytene–diplotene (Fig. 2b). Sequencing RT-PCR products from WT and BW233 (*Hvst1*) anthers revealed that *HvST1* and *Hvst1* produced mRNA encoding 330 and 298 amino acid proteins respectively, confirming the introduction of a premature termination codon in the latter (Fig. 2c and Fig. S4). Screening a further four independent semi-sterile desynaptic mutants (Fig. S5) originating from different cultivars (Betzes, Klages) revealed that they also carried spontaneous mutations in the same microsatellite; BW240 (*des4.af;* from cv Klages), BW241 (*des4.a,* from cv Betzes) and BW229 (*des1.v,* from cv Freja) contain the same 1bp insertion, while BW242 (*des4.h,* from cv Betzes) has a 1bp deletion (Fig. S6). We generated CRISPR/Cas9 knockouts of this gene in *cv* Golden Promise which also caused semi-sterility (Fig. S7).

**Figure 2:**
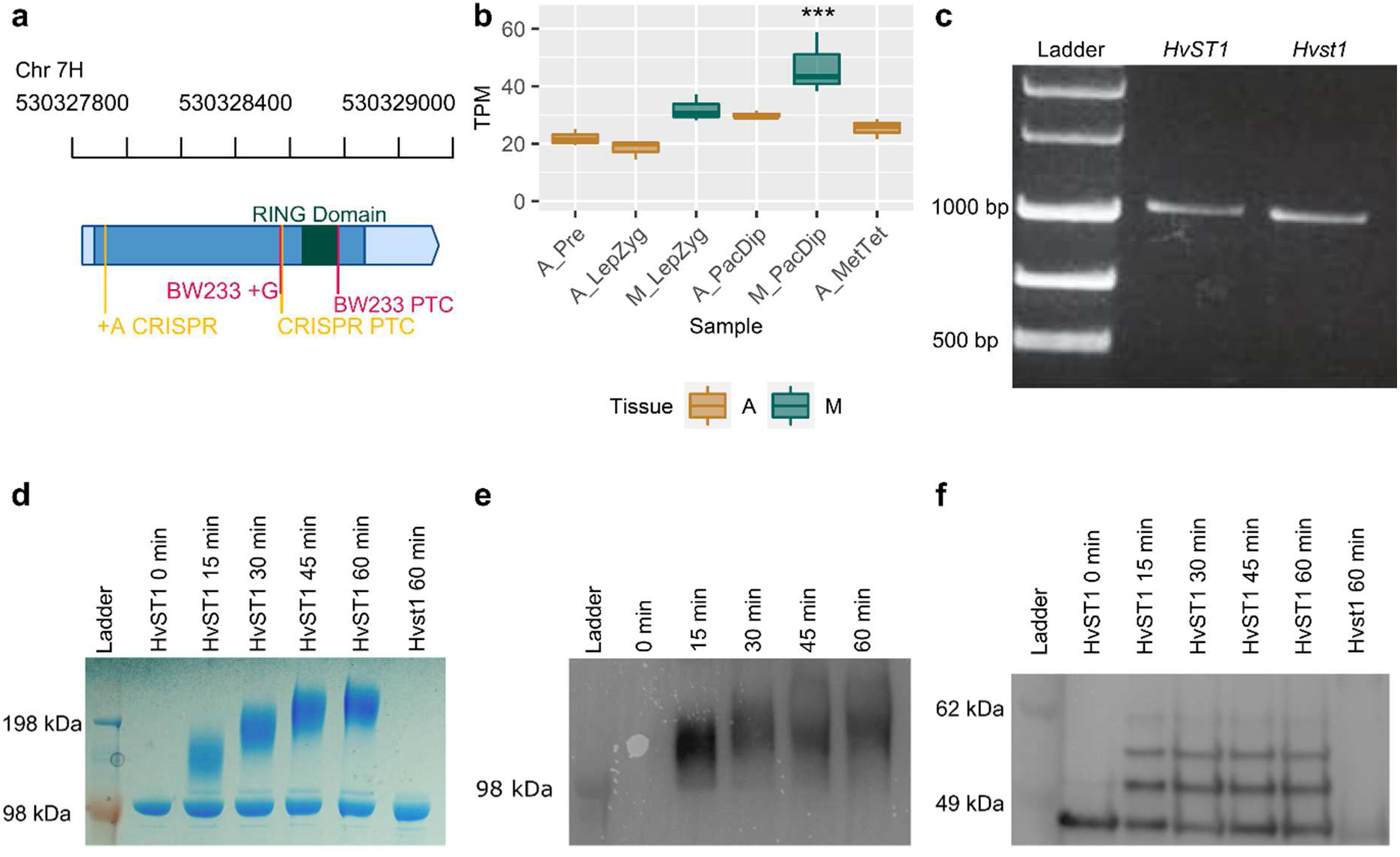
HvST1 characterisation. a) *HvST1* gene model derived from Barley Reference Transcriptome 2 (BaRT2v18; Coulter *et al.*, 2022) RNA sequences mapped to the barley cv Barke genome with the BW233 (*Hvst1)* insertion (+G) and premature termination codon (PTC) indicated in dark pink and the CRISPR/Cas9 insertion and PTC indicated in yellow. The region encoding the RING domain is indicated in dark green. WT 3’ and 5’ untranslated regions (UTRs) are indicated in light blue while WT coding sequences are indicated in dark blue. The physical genomic coordinates are indicated above. b) Expression profile of *HvST1* in time resolved anther and meiocyte expression data (Barakate *et al.*, 2021) where A=Anther, M=Meiocyte, Pre=pre-meiosis, LepZyg=Leptotene–Zygotene, PacDip=Pachytene–Diplotene, and MetTet=Metaphase–Tetrad. The statistical significance (p<0.001) of *HvST1* expression in meiocytes at Pachytene-Diplotene as determined by ANOVA and Tukey’s honest significant difference is indicated above the box. c) agarose gel of FL cDNA from RT-PCR of *HvST1* from Bowman and *Hvst1* from BW233 anthers. d) Coomassie stained SDS-PAGE of autoubiquitination time-course assay. e) anti-ubiquitin conjugate probed western blot of HvST1 autoubiquitination time course assay. f) anti-HvST1 probed western blot of autoubiquitination time course assay.

### HvST1 is a grass-specific functional ubiquitin Ligase

Clustering and phylogenetic analysis of HvST1 places it within a group of grass-specific proteins (Fig. S8), that includes the rice orthologue CYTOSOL LOCALISED RING FINGER PROTEIN 1 (Park *et al.*, 2019) (OsCLR1; Os06g0633500). More recently, Ren *et al*. (2021) reported a study of DESYNAPSIS1 (DSNP1), an induced mutation by irradiation that introduced a 2bp deletion in the exonic region of LOC_Os06 g4270, causing a large reduction of crossovers which was already visible at metaphase and sterility (Ren *et al.*, 2021). Alignment of OsCLR1 and DSNP1 revealed that they are identical. Within grasses HvST1/OsCLR1-like proteins shows strong C-terminal conservation—including the RING domain—with a more variable N-terminal region (Fig. S9). Outside of the Poaceae alignment is limited almost entirely to the E2 interacting RING domain. *OsCLR1* is highly expressed under salt and drought stress (Park *et al.*, 2019). However, this is not a trait shared by orthologues in sorghum, maize, and wheat (Park *et al.*, 2019), nor is *HvST1* differentially expressed under these conditions (Fig. S10), suggesting a unique gain of function in rice OsCLR1/DSNP1.

RING domain containing proteins comprise the largest group of E3 ubiquitin ligases, which confer substrate specificity to the ubiquitination cascade (Dove *et al.*, 2016). The RING domain allows the E3 ligase to recruit E2 conjugating enzymes allowing the transfer of ubiquitin from the E2 to E3-bound protein substrates (Dove *et al.*, 2016; Iconomou and Saunders., 2016). Loss of this domain in *Hvst1* should therefore prevent interaction with E2 conjugating enzymes. In autoubiquitination time course assays, purified HvST1 interacted with the human E1 activating (UBE1) and E2 conjugating enzyme (UBE2D4), producing high molecular weight polyubiquitinated substrates visible via both Coomassie gel staining (Fig. 2d) and western blot with anti-ubiquitin conjugate antibodies (Fig. 2e). Purified *Hvst1* mutant protein did not interact with any E2 (Fig. 2d). Western blotting with anti-HvST1 antibodies confirmed that HvST1 gained mass over the autoubiquitination time course (Fig. 2f). The identity of all protein bands in this assay was further confirmed by mass spectrometry (Fig. S11), confirming the specificity of the HvST1 antibody. Purified HvST1 protein interacted strongly with all human E2 conjugating enzymes in the UBE2D and UBE2E families (Fig. S12), and to a lesser extent with UBE2A, UBE2B, and UBE2N/V1. We conclude that HvST1 is a functional E3 ubiquitin ligase and that loss of the RING domain in *Hvst1* leads to loss of E3 ligase activity.

### *HvST1* mutants exhibit abnormal synapsis

As *Hvst1* mutants are semi-fertile, and display abnormal chromosome segregation (Fig.1), we used Structured Illumination Microscopy (SIM) to compare synapsis progression between the wild type (WT) and *Hvst1* mutants, using antibodies raised against axial (ASY1) and central element (ZYP1) proteins of the synaptonemal complex (SC) (Colas *et al.*, 2017). Axis formation and the initiation of synapsis during early-zygotene were comparable in WT (Fig. 3a) and *Hvst1* (Fig. 3e). By mid-zygotene in the WT most of the chromosomes are paired (Fig. 3b) and the typical tri-partite structure of the SC is visible in the paired region (Philips *et al.*, 2012). In contrast, *Hvst1* meiocytes at the same stage exhibit faltering development with ZYP1 forming abnormal telomeric polycomplex-like clusters (Fig. 3f). At pachytene stage, WT chromosomes are fully synapsed (Fig. 3c) but in *Hvst1* complete synapsis is compromised (Fig. 3g). During diplotene WT chromosomes display a normal tinsel configuration (Colas *et al.*, 2017) as the SC is dissolved (Fig. 3d). However, in *Hvst1*, ASY1 unloading from the SC is perturbed and less organized with apparent misalignment and unpaired regions (Fig. 3h and Fig. S13). The same synapsis defect was observed in insertion mutants BW229, BW240, and knockout Hv*st1^CRISPR/Cas9^*, confirming cytologically that loss of E3 ligase activity in *Hvst1* causes the ZYP1 polycomplex-like phenotype, independently from the original background (Fig. S13 and S14).

**Figure 3:**
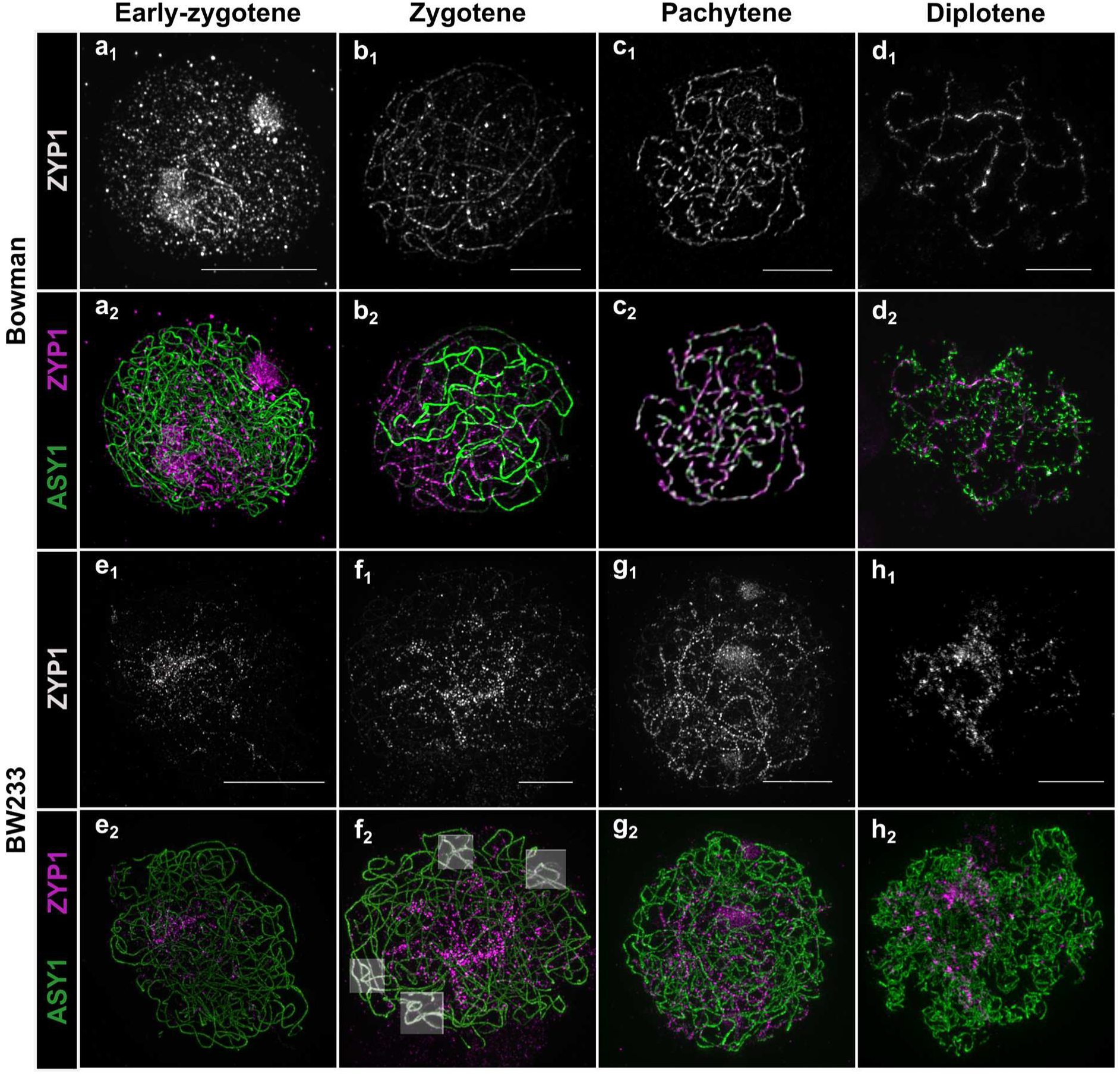
Synapsis in Bowman and BW233. (a-d) Normal synapsis progression in Bowman. a) ZYP1 polymerization starts at one side of the nucleus and elongates during b) zygotene stage. Homologous chromosomes are fully aligned *via* ZYP1 at c) pachytene and get separated at d) diplotene exhibiting normal tinsel chromosomes. e) Somewhat normal initiation of synapsis in BW233. f) zygotene cells of BW233 show abnormal elongation of the ZYP1 “cluster” and unresolved interlocks (highlighted boxes). g) BW233 pachytene-like cell showing persistent HvZYP1 “clustering”. h) BW233 diplotene cells exhibiting abnormal tinsel chromosomes. (a-h) ASY1: green, ZYP1: grey or magenta, scale bars 5 µm.

We found that *Hvst1* mutants had a large number of chromosomal interlocks compared to the WT (Fig. 3f_2_, highlighted boxes) which could be the result of the synapsis delay or abnormal telomere organization. Therefore, we looked at telomere dynamics alongside synapsis and, in both genotypes, the telomeres cluster to one side of the nucleus (Fig.4a, d), suggesting that telomere clustering is not compromised in the mutant. As synapsis progresses, the telomeres move around the nuclear envelope in the WT (Fig. 4b, c) but retain some polarization in *Hvst1* (Fig. 4e, f) indicating that compromised ZYP1 elongation and/or the presence of multiple interlocks affects their movement. The absence of HvST1 function does not therefore compromise initial chromosome alignment but does compromise both progression and dissolution of synapsis.

**Figure 4.**
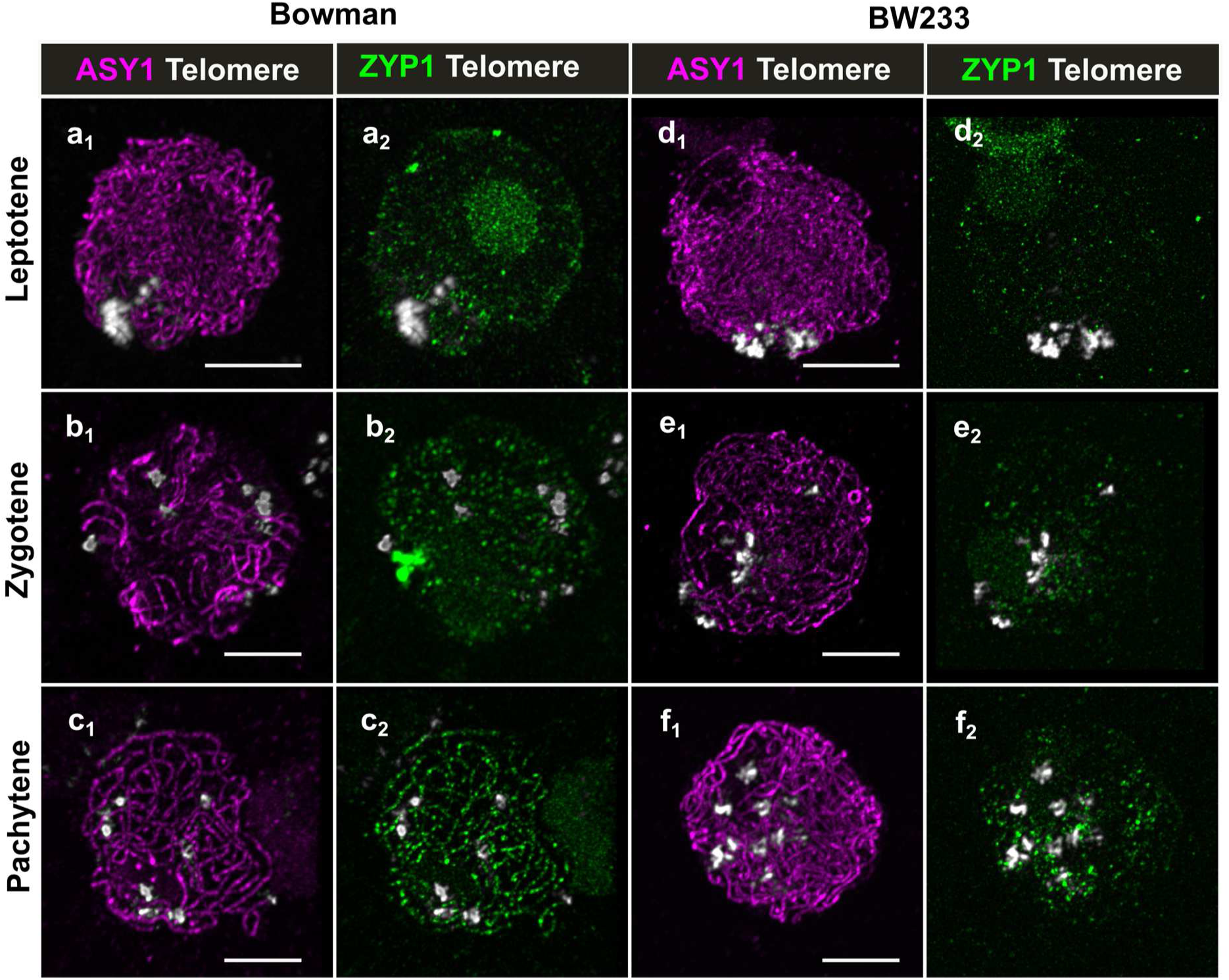
Telomere organization in Bowman and BW233. At leptotene stage, telomeres (grey) cluster at one side of the nucleus in both a) Bowman and d) BW233. As synapsis progresses from b) zygotene to c) pachytene stage, telomeres start to move around the nuclear periphery in Bowman. However, in BW233 e) zygotene and f) pachytene-like stage cells, they are still located at one side of the nucleus. Scale bar 5 µm

We followed synapsis progression and telomere movement using antibodies against ZYP1 and H3K27me3 respectively (Fig.5). At early-zygotene, ZYP1 starts to polymerize in both WT and BW233 and H3K37me3 signals are diffuse (Fig.5a, b). At pachytene, H3K27me3 forms a clear pattern in the WT as described in Baker *et al*. (2015) (Fig. 5e), but not in BW233 where H3K27me3 remains diffuse (Fig. 5d) suggesting a potential change in chromatin state in the mutant. To confirm this, we profiled histone methylation in anthers at pre-meiosis, leptotene/zygotene, pachytene/diplotene and a non-meiotic sample (root). We identified histones H1, H2.A, H2.B, H3, H3 centromeric, and H4 in all samples as well as methylation in H2A, H2B, H3 and H4 (Fig. S15, Table S6). No significant quantitative difference in histone methylation (Kruskal-Wallis test, Benjamini Hochberg correction) was observed (Fig. S15) between WT Bowman and BW233.

**Figure 5.**
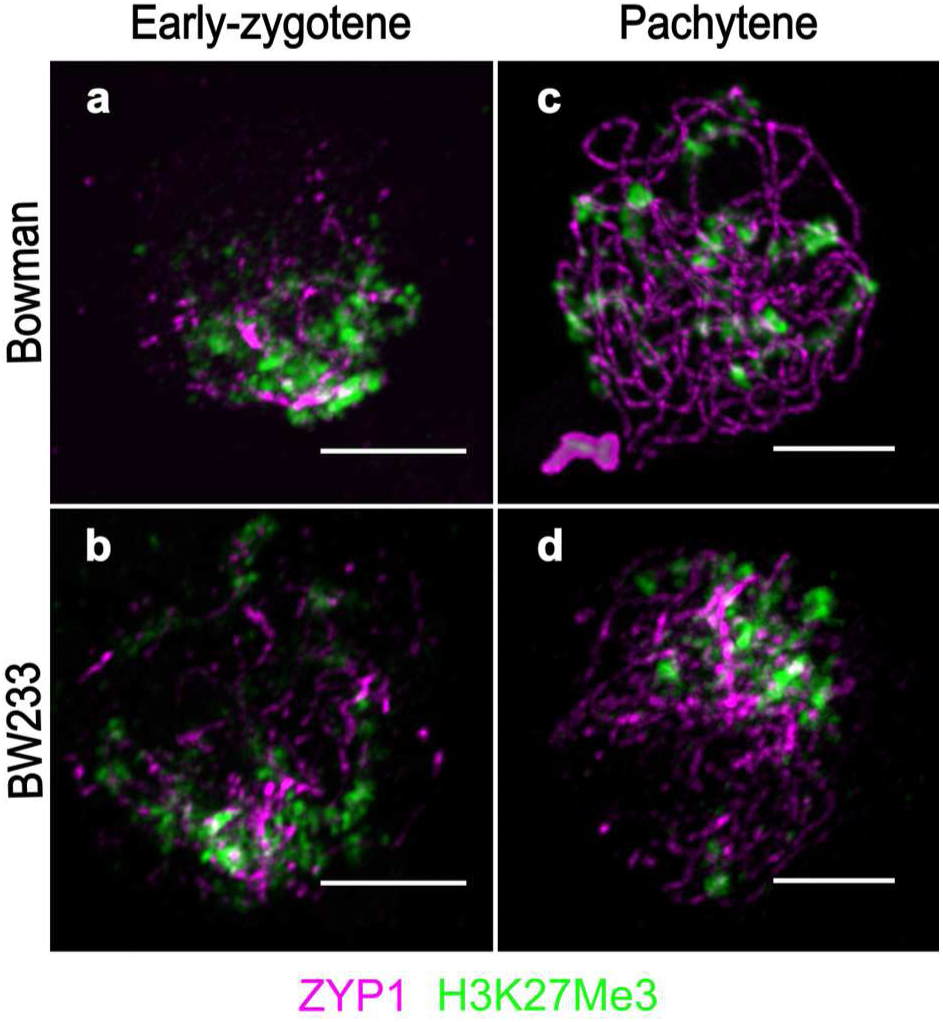
Chromatin behaviour and histone methylation in Bowman and BW233. At the beginning of ZYP1 (magenta) polymerization chromatin seems loose as shown by the diffuse H3K27me3 (green) around the telomere region in both a) Bowman and b) BW233. c) in Bowman at pachytene, the H3K27me3 signal is orderly allowing identification the telomeric regions. d) In BW233, the H3K27me3 signal is more diffuse as a result of abnormal synapsis. Scale bar 5um.

### Recombination is increased in *Hvst1* mutant progeny

To understand the effect of *Hvst1* on recombinaiton we first used antibodies raised against early and late recombination markers. Co-immunolocalisation of HvDMC1 (early recombination) and HvASY1 (chromosomes axes) in both WT and mutant (Colas *et al.*, 2019) showed that HvDMC1 behaviour was similar in both genotypes at the beginning of synapsis (Fig. 6a, 6e) but that HvDMC1 foci persist in the mutant compared to the WT (Fig. 6e) at later stages. However, DMC1 foci counts did not show a significant difference at the leptotene-zygotene transition or at zygotene (Fig. 6i; Student’s T-test, p=0.31 and 0.58), suggesting that HvST1 is not involved in the recruitment of HvDMC1. However, the persistance of foci at later stages is indicative of the delay in synapsis. Co-immunolocalisation of MLH3 (late recombination) and ZYP1 (chromosomes axes) in both WT and *Hvst1* was difficult to conduct due to abnornal synapsis in the mutant. Given the inability to accurately stage late prophase I in BW233 meiocytes due to the perturbation of SC dissolution we compared anthers of similar size as anther size is strongly correlated with meiotic progression (Arrieta *et al.*, 2020). We found that for similarly sized anthers, WT Bowman displayed resolved MLH3 foci at pachytene while BW233 did not (Fig. S16), which is consistent with a delay in crossover resolution (Colas *et al.*, 2016). Co-immunolocalisation of MLH1 (Fig. S17) revealed a similar pattern to MLH3, with a large number of intermediates which persist at later stages. These intermediates converge into larger stretches as prophase I progresses but fail to resolve as discrete foci (Fig. S7f). The class I crossover resolution marker HEI10 (Ziolkowski *et al*., 2017; Serra *et al.*, 2018) typically loads on chromosome axes at zygotene (Fig. 6b) and co-localises with ZYP1 at pachytene (Fig. 6d) in WT Bowman. In BW233, although HEI10 maintains co-localisation with ZYP1, it does not form typical distinct foci but co-localises with the abnormal ZYP1 polycomplex-like structure (Fig. 6f, h).

**Figure 6:**
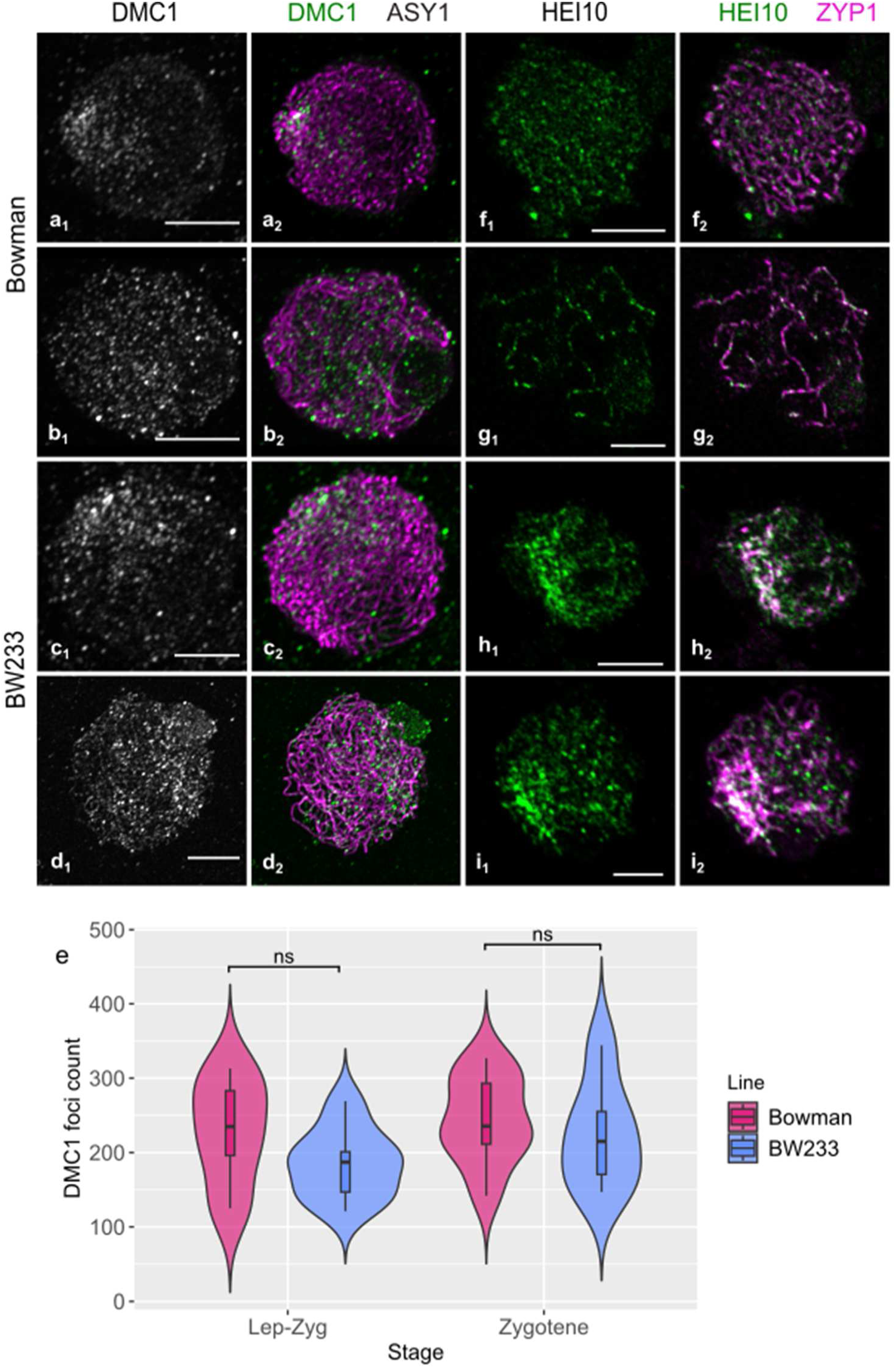
DMC1 and HEI10 behaviour in Bowman and BW233. At leptotene stage, DMC1 foci (green, grey) cluster at one side of the nucleus in a) Bowman and disperse along the ASY1 axes (Magenta) during b) zygotene. In BW233, we see the same behaviour at c) leptotene and d) zygotene. e) Violin plot of DMC1 foci count showing no significant difference between Bowman and BW233. HEI10 foci load on ZYP1 axes in both f,g) Bowman and h,i) BW233, but they resolve as large foci in g) bowman pachytene cells while they cluster around the ZYP1 polycomplex in h,i) BW233. scale bar 5µm

Due to the stickiness of *Hvst1* chromosomes, the unclear MLH3 labelling and the HEI10 polycomplex, it was not possible to obtain accurate chiasma or MLH3/HEI10 foci counts from immunocytology of meiocytes. We therefore directly assessed recombination in *Hvst1* mutant progeny by genetic analysis of 400 F_3_ plants derived from 24 F_2_ families selected from a cross between BW233 (*Hvst1*) x cv. Barke (*HvST1*) after using Marker Assisted Selection (MAS) to identify genotypes homozygous for *HvST1* or *Hvst1.* We focused initially on chromosomes 1H, 5H, and 6H using 48 KASPar markers (LGC Genomics). We observed that despite indicative observations in metaphase I meiotcytes the result of *Hvst1* mutation was a significant increase in distal recombination events in *Hvst1* compared to the WT F3 families for all three chromosomes. To confirm that increased recombination in *Hvst1* was genome-wide, we then chose 95 of the 400 F_3_ plants and genotyped them using the barley 50K iSelect SNP array (Bayer *et al.*, 2017). We calculated genome wide recombination frequency after filtering for poorly mapped markers and F_2_ recombination events (Fig. S18). Total observed crossovers increased in *Hvst1* compared to *HvST1* across all chromosomes—primarily distally—by an average of 76% and up to 149% (Fig. 7; Table S7). This shows that the absence of HvST1 E3 ligase activity increased meiotic recombination in those meiocytes which formed viable gametes.

**Figure 7:**
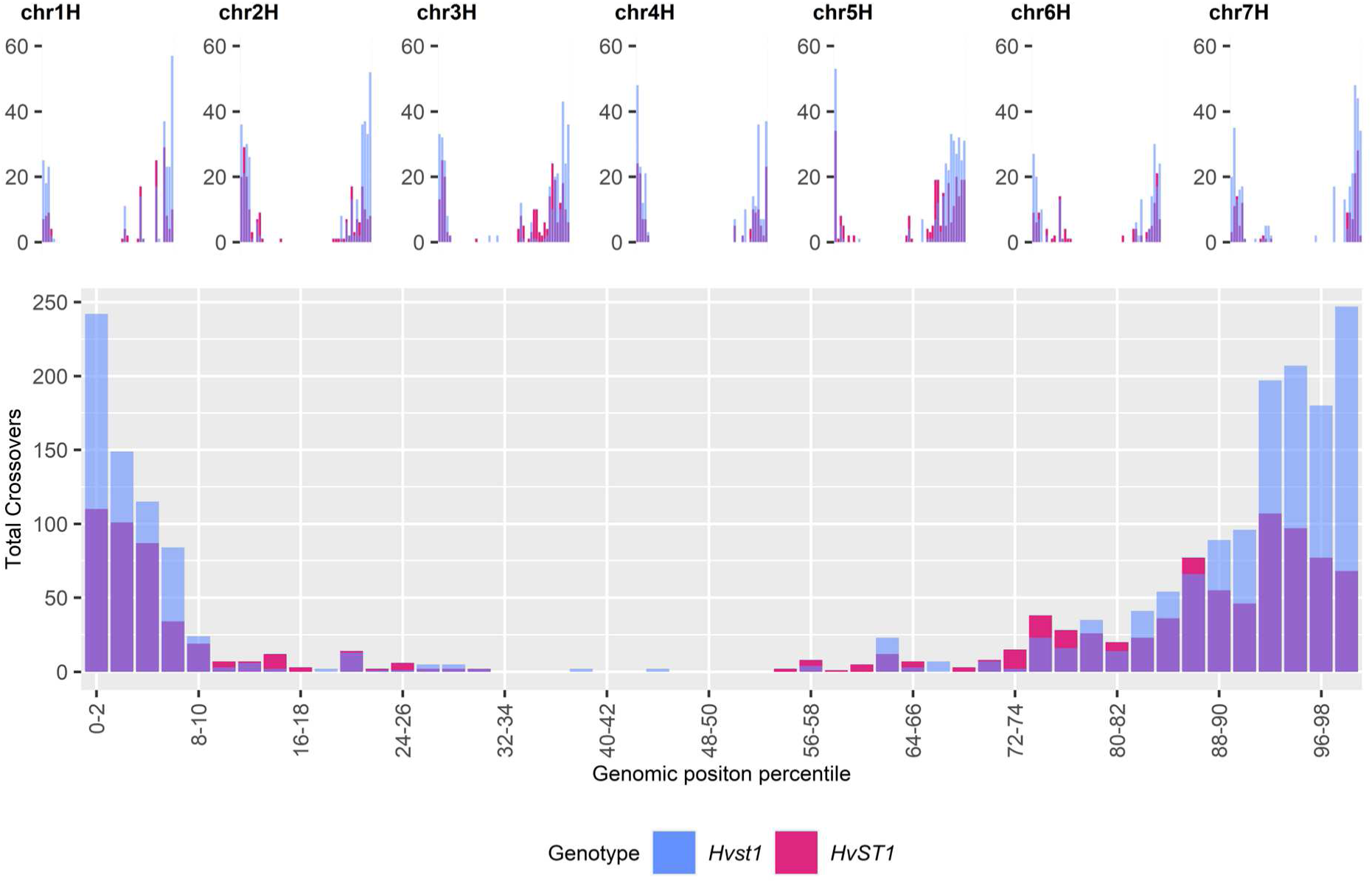
Recombination. Comparison of the total number of crossover events detected using 50K iSelect markers across all seven chromosomes (represented individually above and combined below) between *Hvst1* (in Blue; n=95) and *HvST1* (WT; in pink; n=95) per binned (2%) genomic position (Mbp) percentile.

### HvST1 is capable of ubiquitinating ASY1

To determine if abnormal synapsis observed in *Hvst1* mutants might be explained by failure of SC protein ubiquitination (required for their removal by the proteasome), we conducted an *in vitro* substrate ubiquitination assay with purified HvASY1 and HvST1 proteins. We found that HvASY1 is ubiquitinated in the presence of HvST1 and all other required components of the ubiquitination cascade (Fig. 8a), visible as an increase in mass on western blots probed using anti-ASY1 antibody. No high molecular weight protein is labelled in anti-ASY1 western blots in the absence ATP, with purified Hvst1, or in the absence of HvASY1 (Fig. 8a) suggesting that the observed ubiquitination of HvASY1 is dependent on the HvST1 RING domain and the ubiquitination cascade. To confirm that high molecular weight HvASY1 was ubiquitinated, the polyubiquitinated products of this assay were captured using GST-TUBE and treated with broad spectrum deubiquitinating enzyme which recovered HvASY1 at its original mass (Fig. 8b).

**Figure 8.**
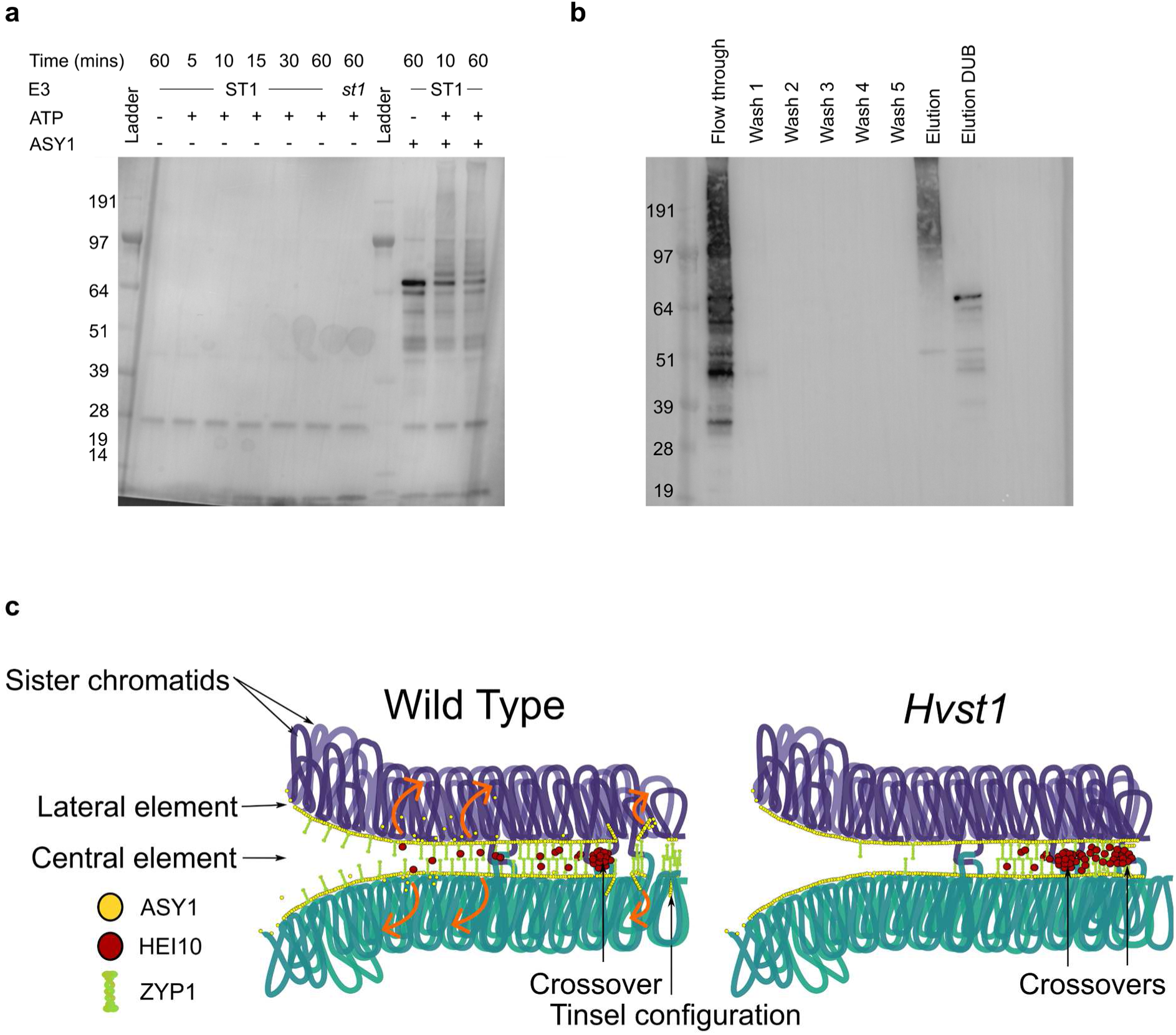
ASY1 ubiquitination and Model. a) Anti-ASY1 western blot of HvST1 autoubiquitination time course and HvST1-ASY1 substrate ubiquitination assay; lane description from left: 1-ladder; 2-HvST1 autoubiquitination no ATP control, 3-7: HvST1 autoubiquitination time course, 8: Hvst1 autoubiquitination control, 9: ladder, 10: HvST1-ASY1 substrate ubiquitination no ATP control, 11-12: HvST1-ASY1 substrate ubiquitination assay. b) Western blot of fractions recovered from GST-TUBE capture of polyubiquitinated products of HvST1-HvASY1 substrate ubiquitination assay probed with anti-HvASY1 antibody. HvASY1 antibody labels TUBE captured ubiquitinated protein in the eluted fraction and upon treatment of this fraction with broad spectrum deubiquitinating (DUB) enzyme ASY1 is returned to its original size (∼64 KDa). Marker sizes in KDa are indicated on the left. c) proposed model of the role of HvST1 in synapsis (purple loops= homologue 1; sea green loops=homologue 2. In the wild type, synapsis is synchronized and ZYP1 polymerises between the two homologues dependent on continuous ASY1 turnover while the pro-crossover factor HEI10 diffuses along the ZYP1 axes. In *Hvst1* mutants, ASY1 turnover is compromised leading to partial synapsis and formation of the ZYP1 cluster while diffusion of HEI10 is constrained to partially synapsed regions, affecting coarsening dynamics by increasing the subtelomeric concentration of HEI10, leading to more crossovers which are closer together in this region.

## Discussion

In plants, mammals, and budding yeast the formation of crossovers and synapsis are tightly linked (Grey *et al.*, 2022). We have previously shown that mutations in barley meiotic genes dramatically affect meiosis and, in general, reduce recombination (Colas *et al.*, 2016: Colas *et al.*, 2019) but also that mutations in conserved meiotic genes in barley (Colas *et al.*, 2016; Colas *et al.*, 2023) do not always lead to the expected phenotype when compared to other plants, including *Arabidopsis* (Jackson *et al.*, 2006; Bleuyard *et al.*, 2004) or rice (Ren *et al.*, 2021).

Here, we have identified and characterized an E3 ubiquitin ligase that is highly expressed during meiosis, providing the first evidence of a unique ubiquitination pathway that regulates meiosis in barley. We have called this gene *HvST1* (Sticky Telomeres 1), due to the stickiness of chromosomes at metaphase I that seemed to be in the telomere region. This stickiness made chiasma counts in *Hvst1* challenging but we did find a significant increase in the average number rod bivalent chromosomes and decrease in ring bivalent chromosomes were observed in *Hvst1* metaphase compared to the WT. We also observed a large variation in chiasma number in BW233 compared to the WT which in return led to a significant, albeit small, decrease of chiasma in BW233 compared to the WT, reflecting different metaphase configurations.

The most significant phenotype of the *Hvst1* genotype was abnormal synapsis due to the formation of ZYP1 polycomplex-like “clusters” near the telomere region. Contrary to our initial hypothesis, we found that initial formation of the telomere cluster was not altered in the mutant but that the telomeres tended to remain located at one side of the nucleus alongside ZYP1 clustering at the onset of delayed synapsis. Moreover, analysis of the chromatin state in BW233 using antibodies targeting methylated histones indicated altered chromatin state, although proteomic analysis of histone methylation did not show statistically significant differences in any such modification. This suggests that HvST1 activity does not regulate the relative amount of histone methylation although it is still possible that loss of HvST1 function may affect the distribution of histone methylation along the chromatin length. Accordingly, the apparent reduction of chromatin compaction in BW233 is likely a reflection of delayed synapsis.

We also found that altered distribution of ZYP1 in *Hvst1* constrained the distribution of HEI10 in late prophase, while HvDMC1 and MLH3 recruitment was unaltered, but failed to resolve into discrete foci. The failure of MLH3 foci formation and concentration of HEI10 in the sub telomeric polycomplex also prohibited cytological counting of class I crossovers based on recombination intermediate foci in late prophase I. However, F3 50K genetic recombination data showed a clear and substantial increase in the number of sub-telomeric recombination events in *Hvst1* relative to the WT. Further work is required to determine the pathway responsible for this increase in recombination. Successful progeny included in the genetic recombination data likely represent a bias towards less extreme male meiotic phenotypes exhibiting better chromosome alignment (Fig. S9) and proper segregation as these are more likely to develop to form fertile pollen. As such, the genetic recombination data, while a direct reflection of the effective recombination rate between generations, is an incomplete representation of cytological CO repair pathways. An increase in recombination alongside a higher number of rod bivalent chromosomes at metaphase can be explained by a loss of obligate crossovers alongside crossovers which occur closer together which can be counted as a single chiasma as reported in the wheat *fancm* mutant Desjardins *et al.*, 2022) and the *Arabidopsis zyp1* mutants (France *et al.*, 2021). The phenotype of *Hvst1* bears some other similarities to *zyp1* null mutants in *Arabidopsis* including persistent chromosome interlocks and failure of ASY1 depletion (Capilla-Perez *et al.*, 2021; France *et al.*, 2021). This indicates that much of the *HvST1* phenotype might derive from downstream effects of the failure or severe delay in complete ZYP1 polymerization.

Polyubiquitination of HvASY1 by HvST1 but not Hvst1 *in vitro* provides an indication that the lack of ZYP1 polymerization observed in *Hvst1* may result from the loss of this post translational modification of ASY1. This observation supports the findings of Osman, *et al*. (2018) who previously identified proteasomal and ubiquitination related proteins in association with ASY1 in a pulldown from *Brassica oleracea* and is consistent with the observed localisation of ubiquitination to the chromosomal axes during meiotic prophase (Rao *et al.*, 2017; Orr *et al.*, 2021b). RING type E3 ubiquitin ligases confer specificity to the ubiquitination cascade by binding both E2-conjugating enzymes, thereby increasing the reactivity of E2-Ub conjugates, and substrate proteins, catalysing the direct transfer of ubiquitin to the substrate protein itself, or existing polyubiquitin chains on its surface (Dove *et al.*, 2016; Iconomou and Saunders, 2016). The requirement for a functional RING domain for HvST1-HvASY1 interaction is evident in the lack of ASY1 ubiquitination in the absence of the E2 interacting RING domain in Hvst1 (Fig. 8a). The canonical fate of polyubiquitinated proteins is proteasomal degradation, although a range of substrate fates can arise from ubiquitination determined by polyubiquitin chain topology, which is itself largely driven by a preference for particular lysine residues on the part of the interacting E2s (Dove *et al.*, 2016). Combining this observation with the observed retention of ASY1 on the axis in late prophase I in BW233 (Fig. 3g-h), peak of *HvST1* expression in meiocytes at Pachytene-Diplotene (Fig 2b), the co-localisation of pro-crossover factor HEI10 with ZYP1 during delayed BW233 ZYP1 polymerisation (Fig. 6f, 6h), and the significant increase in distal recombination events in homozygous *Hvst1* progeny in both KASP and iSelect 50K analysis (Fig. 7), we propose a model for HvST1 function in barley meiosis (Fig 8c) in which ASY1 turnover in early prophase I is required for ZYP1 polymerisation and by extension for normal SC formation, crossover resolution, and interference. A requirement for lateral element ubiquitination and proteasomal degradation for synapsis progression has been previously demonstrated in mice where interaction of the SKP1-Cullin-F-box (SCF) complex in conjunction with F-box protein FBXO47 with ASY1 orthologue HORMAD1 is required for normal progression of synapsis (Guan *et al.*, 2022; Ma *et al.*, 2024).

In *Hvst1* mutants HEI10 distribution is clearly constrained by altered ZYP1 distribution resulting in a localised increase in HEI10 concentration and long stretches of HEI10 in partially synapsed regions as opposed to discrete foci. It has been demonstrated that the distribution of HEI10 is constrained by ZYP1—so long as ZYP1 is present—and that altered ZYP1 distribution can affect crossover number and distribution (Capilla-Perez *et al.*, 2021; France *et al.*, 2021; Fozard *et al.*, 2023). In this context, the observed increase in distal recombination in *Hvst1* may be best understood within the proposed HEI10 coarsening model (Morgan *et al.*, 2021). It has been demonstrated that an increase in HEI10 concentration results in a reduction in crossover interference and an increase in the total number of crossovers (Ziolkowski *et al.*, 2017). The HEI10 coarsening model proposes that class I crossover number and distribution can largely be attributed to the concentration, SC restricted diffusion, and reduced rate of escape over time of HEI10 from larger foci at recombination intermediates which are then resolved as crossovers. However, altered synapsis in *Hvst1* mutants prohibited accurate cytological counting of class I or class II recombination intermediates (Fig. 6, S16, S17), putting determination of the crossover pathway responsible for the increase of recombination observed in KASP marker and 50K recombination analysis outside the scope of this work.

Recent description of the rice *Osclr1*/*dsnp1* meiotic phenotype highlights several similar immunocytological observations, including the formation of a ZEP1 polycomplex, the ZYP1 equivalent in rice, and failure of PAIR2 depletion, the ASY1 equivalent in rice, indicating conserved meiotic function in grasses (Ren *et al.*, 2021). However, the induced *dsnp1* mutation rendered these plants completely sterile, *Osdsnp1* metaphase spreads do not show the lack of chromosome condensation observed in *Hvst1,* and the ZEP1 polycomplex does not appear to retain subtelomeric localization as in *Hvst1* (Ren *et al., .* 2021). The authors also reported reduction in class I crossovers based on reduced chiasmata counts in *Osdsnp1* compared to the WT and decreased HEI10 foci counts in a *Osdsnp1/zep1* double mutant background when compared to both the WT and *zep1* single mutant (Ren *et al.*, 2021). The induced mutation in *Osdsnp1* is highly similar to *Hvst1* occurring early in the RING domain and presumably resulting in loss of E3 ligase activity (Ren *et al.*, 2021). Although OsDSNP1/OsCLR1 has a reported gain of function in response to heat and drought stress which is not conserved in other grasses (Park *et al*., 2019), indicating some evolutionary divergence, it does not seem likely that the function of this protein is meaningfully divergent from HvST1 in meiosis. The reported reduction in *Osdsnp1* chiasma may reflect the greater proximity of crossovers we observe in genetic recombination data, resulting in inaccurate counts as described by Desjardins *et al*. (2022). The slight reduction of HEI10 foci in the *Osdsnp1/zep1* double mutant background compared to the WT might indicate that the increase in recombination we observe in *Hvst1* is due to class II crossover resolution, or that the altered ZYP1 environment and partial synapsis is essential to the increase in recombination observed in *Hvst1*. The comparatively large reduction in HEI10 foci in *Osdsnp1/zep1* compared to the *zep1* single mutant is intriguing, possibly indicating a role for HvST1 in regulating class I crossover resolution beyond its impact on ZYP1 polymerization, whether through ASY1 ubiquitination or of other targets. Complete sterility and the distinct ZEP1 polycomplex behaviour in *Osdsnp1* most likely reflect fundamental differences in genome size, chromatin organization, and timing during meiosis between rice and barley. Further work is required to identify and validate the targets of HvST1 E3 ligase activity *in vivo* and to elucidate the protein-protein interactions underlying the *Hvst1* meiotic phenotype.

There is increasing interest in methods of altering the recombination landscape in plants through targeting of post-translational modifications and the use of methods such as virus induced gene silencing (VIGS) to downregulate crossover suppressive genes in order to reduce the time, resources, and emissions associated with generating novel crop varieties with desirable traits (Desjardins *et al.*, 2020. Raz *et al.*, 2020). Further investigation of the molecular mechanics of *HvST1* might reveal new pathways and additional targets for such approaches. Similarly, identifying alternative mutations in *HvST1*, its promoter, targets, or proteins regulating its activity might result in a similar increase in recombination without the same degree of semi sterility which would enhance its use in plant breeding. While it is presently unclear exactly how subtelomeric recombination is increased in *Hvst1,* this mutation could find immediate practical application in breeding programs to increase recombination—in particular, between chromosomes of distantly related genotypes that are being exploited as a source of novel traits such as disease, heat, and drought resistance and to disrupt stubborn linkage drag. Serra *et al*. (2018) demonstrated that combining *HEI10* overexpression with mutation of *recq4*, an inhibitor of class II COs, led to an additive increase in recombination in Arabidopsis. Combining *Hvst1* with the recently described *Hvrecql4* mutation (Arrieta *et al.*, 2021) may also lead to such an additive effect and may provide further insight into the pathway leading to the observed increase in recombination in *Hvst1*.

## Acknowledgments

The authors would like to thank Prof. Ron Hay, Dr Federico Pelish, and Dr Frank Wright for providing advice on the experiments and Dr Michael Skelly for providing the GST-TUBE construct. This research was funded by Biological Science Research Council Grant BB/T008636/1, with support data generated under the European Community’s Seventh Framework Programme *FP7/2007-2013 (n*° 222883), the European Research Council Grant Shuffle (Project ID: 669182), and the Scottish Government’s Rural and Environment Science and Analytical Services Division work programme Theme 2 WP2.1 RD1 and RD2. OMX microscopy was supported by the Euro-BioImaging PCS to I.C. and through the MRC Next Generation Optical Microscopy Award (Ref: MR/K015869/1) at the CAST Facility, University of Dundee. All other microscopy work was supported by the Imaging Facility of the James Hutton Institute. SM was supported by the joint James Hutton Studentship program and EASTBIO fellowship (2013-2017). pDEST-HisMBP was a gift from David Waugh (Addgene plasmid #11085).

## Author Contributions

IC, LR, JO, SM and RW designed the experiment. SM, JO, DL, MM, AB, and NM conducted the experiments and data analyses. JO, RW, and IC wrote the manuscript. All authors reviewed the manuscript.

## Data availability

Barley 50K iSelect recombination data is available on FigShare (https://doi.org/10.6084/m9.figshare.21427953.v1). Raw immunocytology images are available from the corresponding author upon reasonable request.

## Code availability

Code used in data analysis and plotting for this study is available at https://github.com/BioJNO/HvST1.

