## supplemental figures for "Loss of E3 ligase *HvST1* function substantially increases recombination"

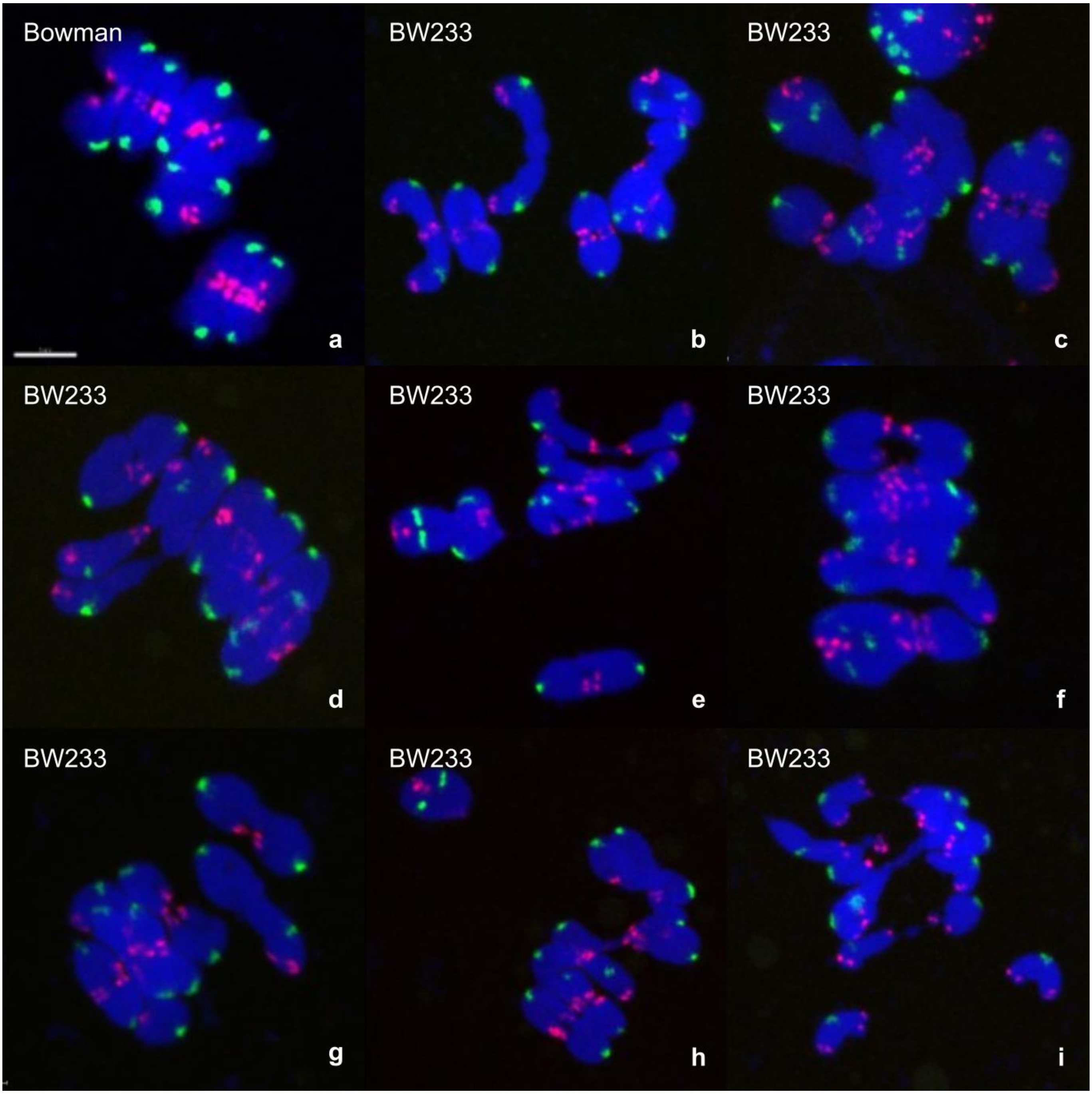


**Figure S1. BW233 Metaphase configuration**

Metaphase spreads of Bowman (*HvST1*) and BW233 (*Hvst1*) with labelled centromeres (in green), telomeres (in red), and chromatin (in blue). a) Bowman metaphase showing 7 ring bivalents with aligned telomeres. b-i) BW233 metaphases showing sticky chromosome configuration predominantly linked from their sub-telomeric regions and a range of ring, rod, and univalent chromosomes. Scale bar 10 µm.

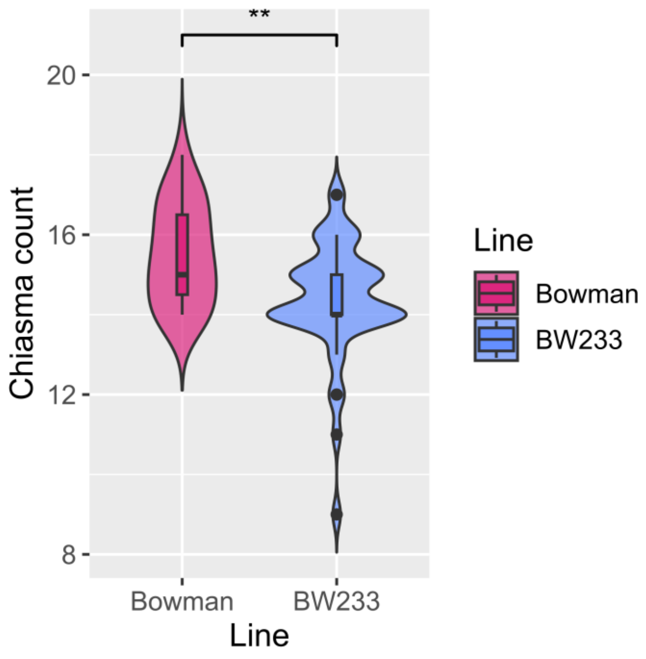


**Figure S2. Chiasma counts from Bowman and BW233 metaphase spreads**
(a) Violin plot of chiasma counts from Bowman (n=19) and BW233 (n=44) meiocytes at metaphase. Statistical significance of the comparison of mean chiasma counts between Bowman and BW233 (Wilcoxon rank sum test, p=0.0055) is indicated above the violins.

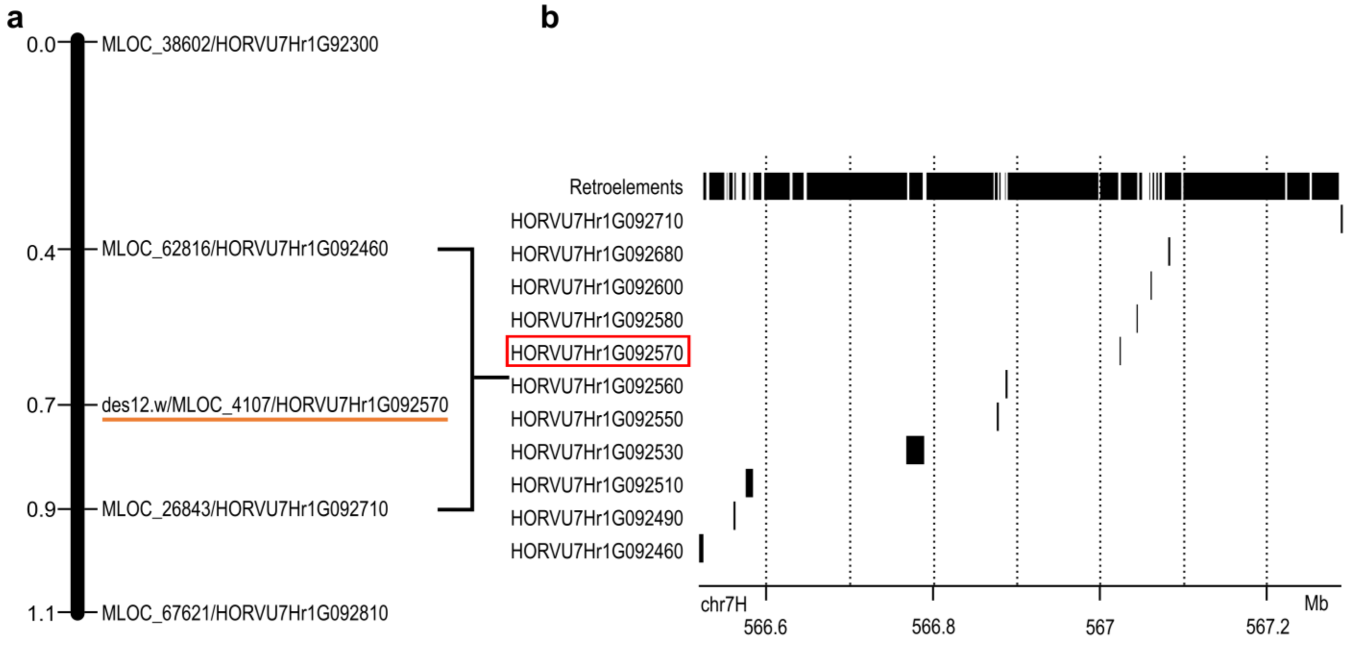


**Figure S3. *des12.w* Initial Mapping**
(a) 1cM *des12.w* interval (b) 0.5cM *des12.w* interval showing 9 high confidence gene models and retrotransposable elements extracted from Mascher et al. (2017). The red box indicates *des12.w (Hvst1)*

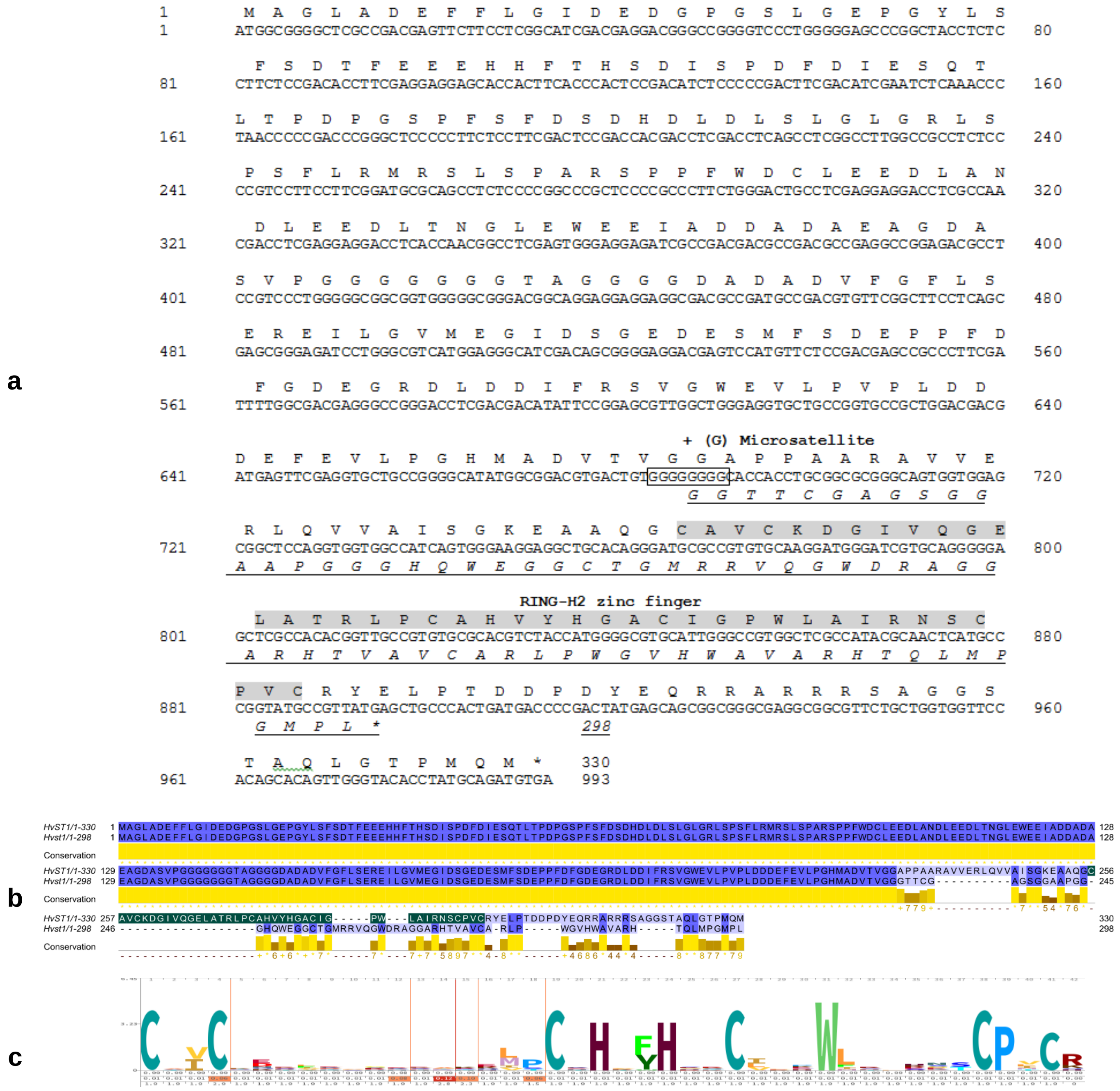


**Figure S4. *des12.w m*utation**

a) *HvST1* gene and protein sequences. b) Grey shading marks the RING-H2 zinc finger domain. c) Precited RING-H2 zinc finger logo with multiple protein alignments of 127 RING-H2 finger proteins from dicots and monocots (Katoh, Misawa, Kuma & Miyata 2002; Katoh & Standley 2016; Wheeler, Clements & Finn 2014).

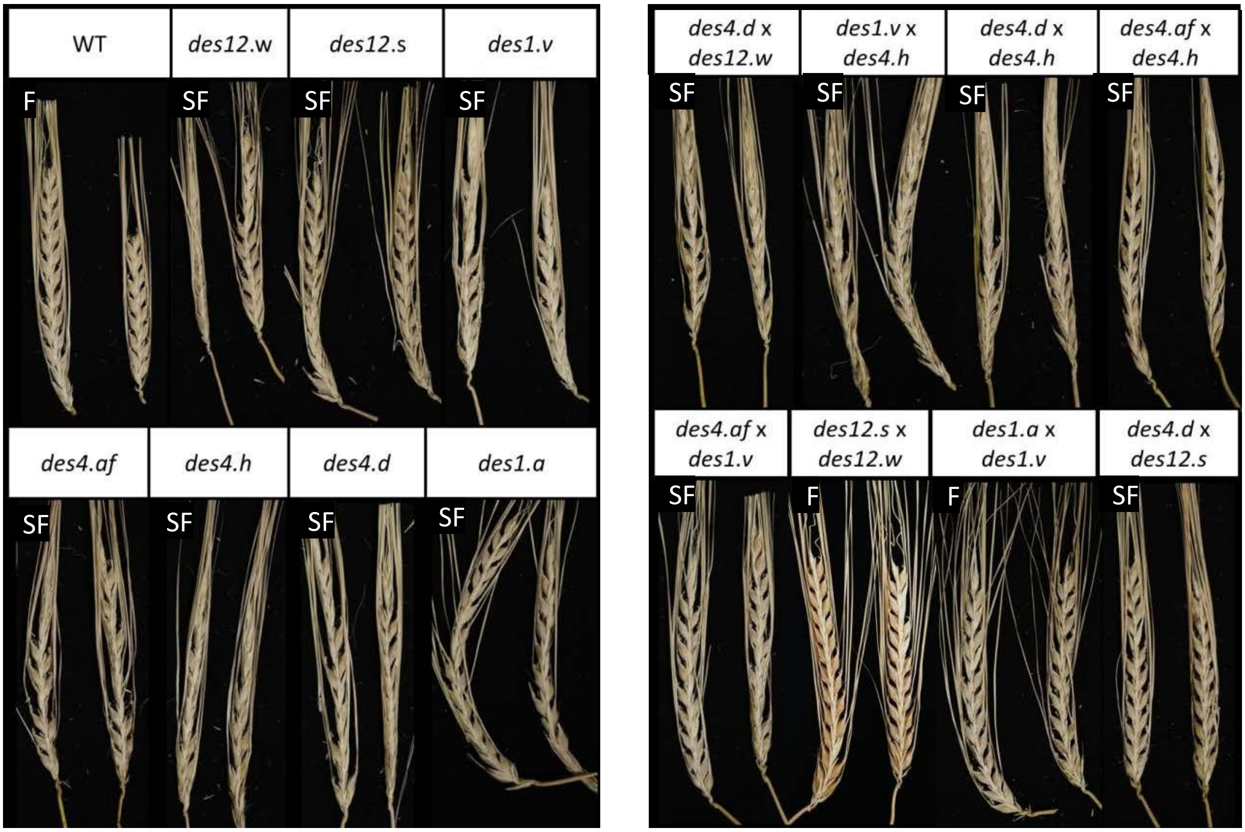


**Figure S5. Fertility of the allelic test crosses and their parental lines**

Ears of Bowman (WT), BW233 (*des12.w*), BW232 (*des12.s*), BW228 (*des1.a*), BW229 (*des1.v),* BW240 (*des4.af*), BW241 (*des4.d*), BW242 (*des4.h*) and *desynaptic* crosses for allelism test. Semi-fertile crosses indicate that the mutants carry the same genes. BW232 (*des12.s*) and BW228 (*des.1a*) are independent mutations in a different genes than the other *desynaptic* mutants. SF: Semi-fertile; F: Fertile.

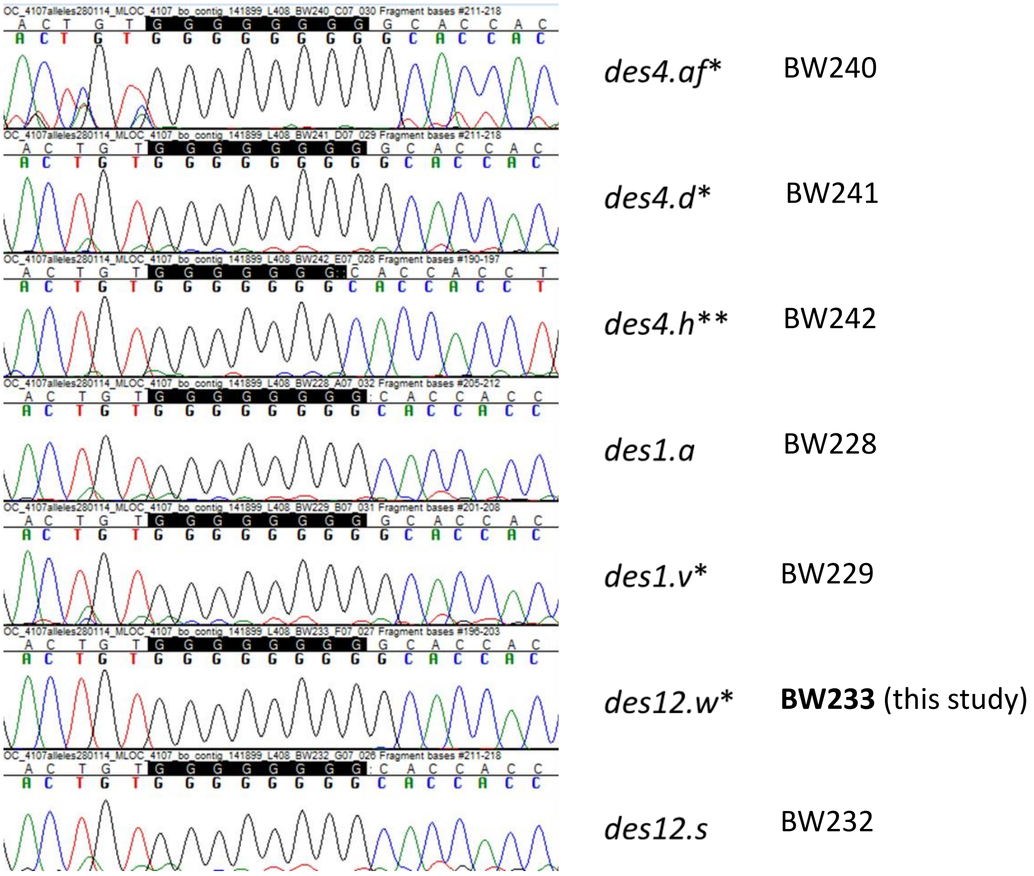


**Figure S6. Clipped Sanger sequencing chromatograms for *des12, des4,* and *des1* alleles.** Confirming the allelism test, BW240*,* BW241*, and* BW229 carry an identical 1bp insertion (*) at the 8 bp small microsatellite, similar to BW233*,* while BW242 contains a 1bp deletion (**) in this same microsatellite. All mutations caused semi-fertility in the original lines and reciprocal crosses.

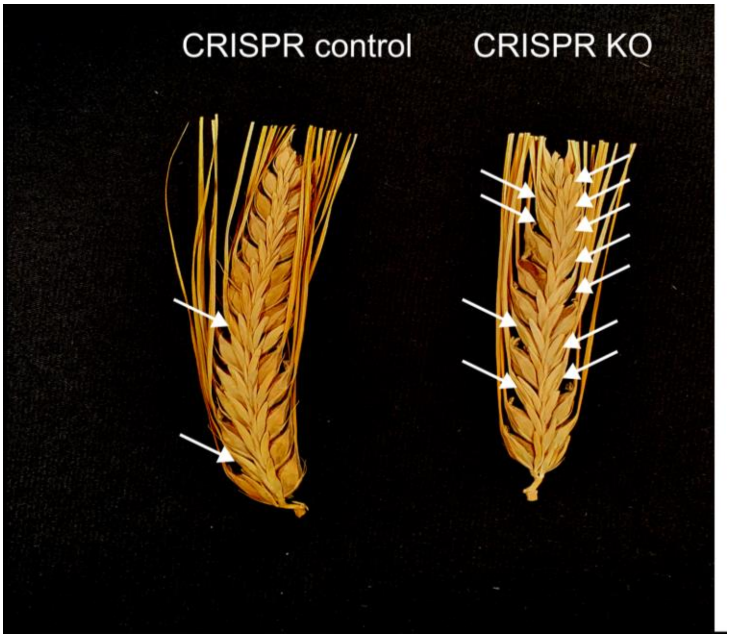


**Figure S7. CRISPR/*cas9* Line for *HvST1***
Knock out of *HvST1* by CRISPR/*cas9* causes semi fertility. Missing seed are indicated by white arrows.

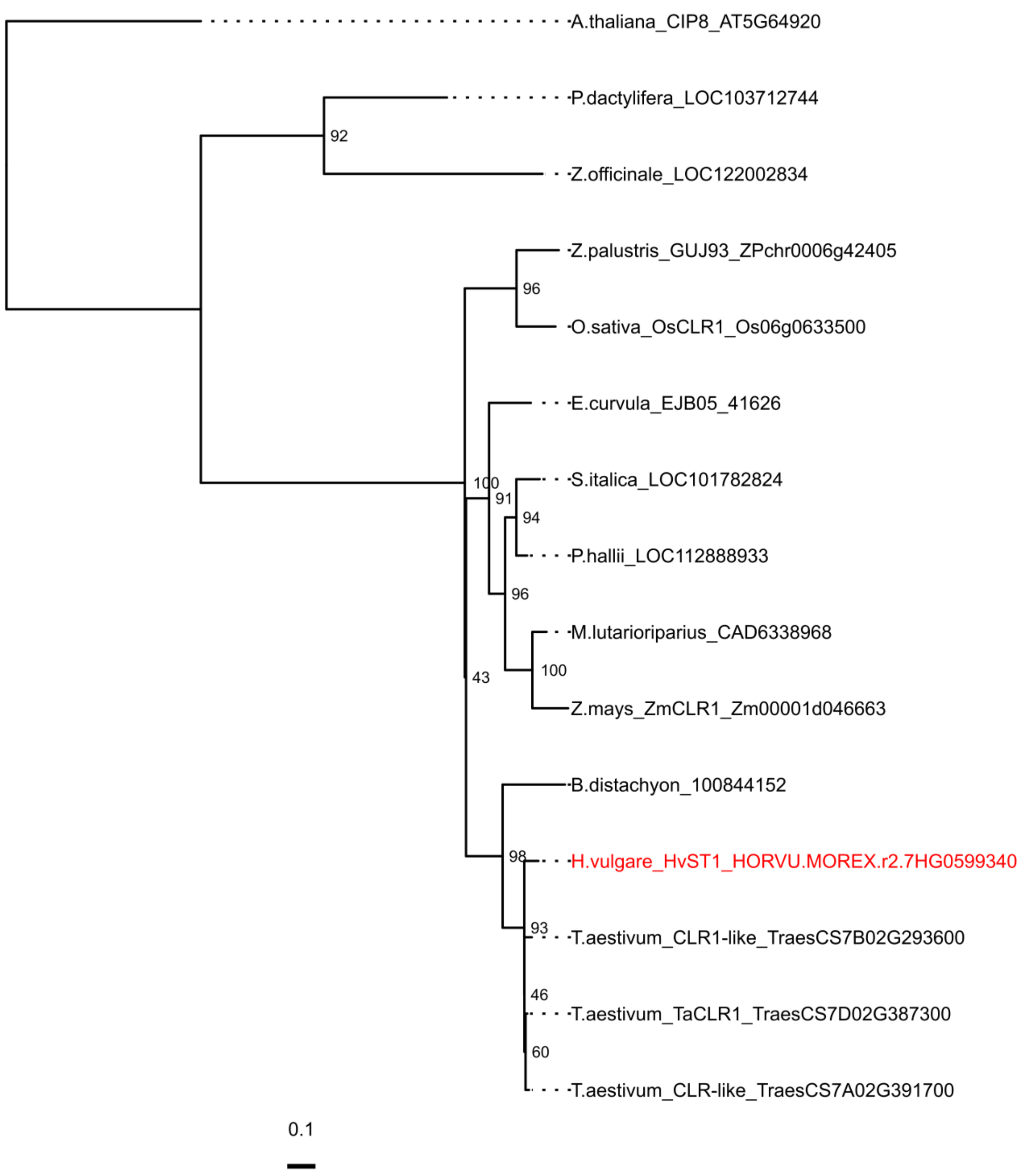


**Figure S8.** Maximum likelihood phylogeny of *HvST1* and its orthologues with *A. thaliana* RING E3 *CIP8* as out group. Nodes are labelled with bootstrap support.

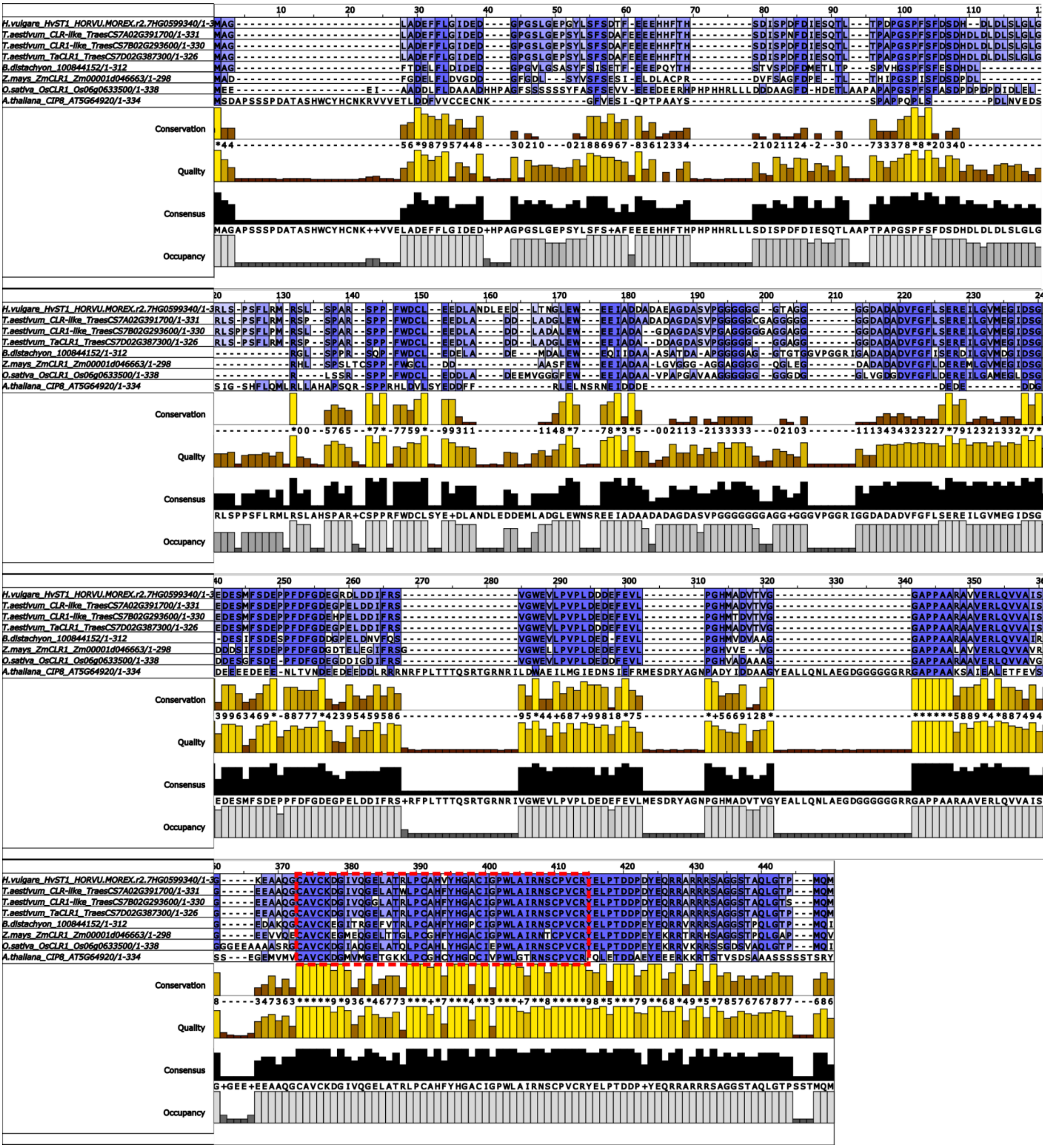


**Figure S9.** Amino acid alignment of *HvST1*, its orthologues in the Poaceae, and *A. thaliana CIP8*. Alignment between orthologues in the Poaceae and *Arabidopsis* is largely limited to the conserved C-terminal RING domain highlighted in the red box. Within the Poceae the C-terminal region is very well conserved with more variation towards the N-terminal region, particularly in *O. sativa.*

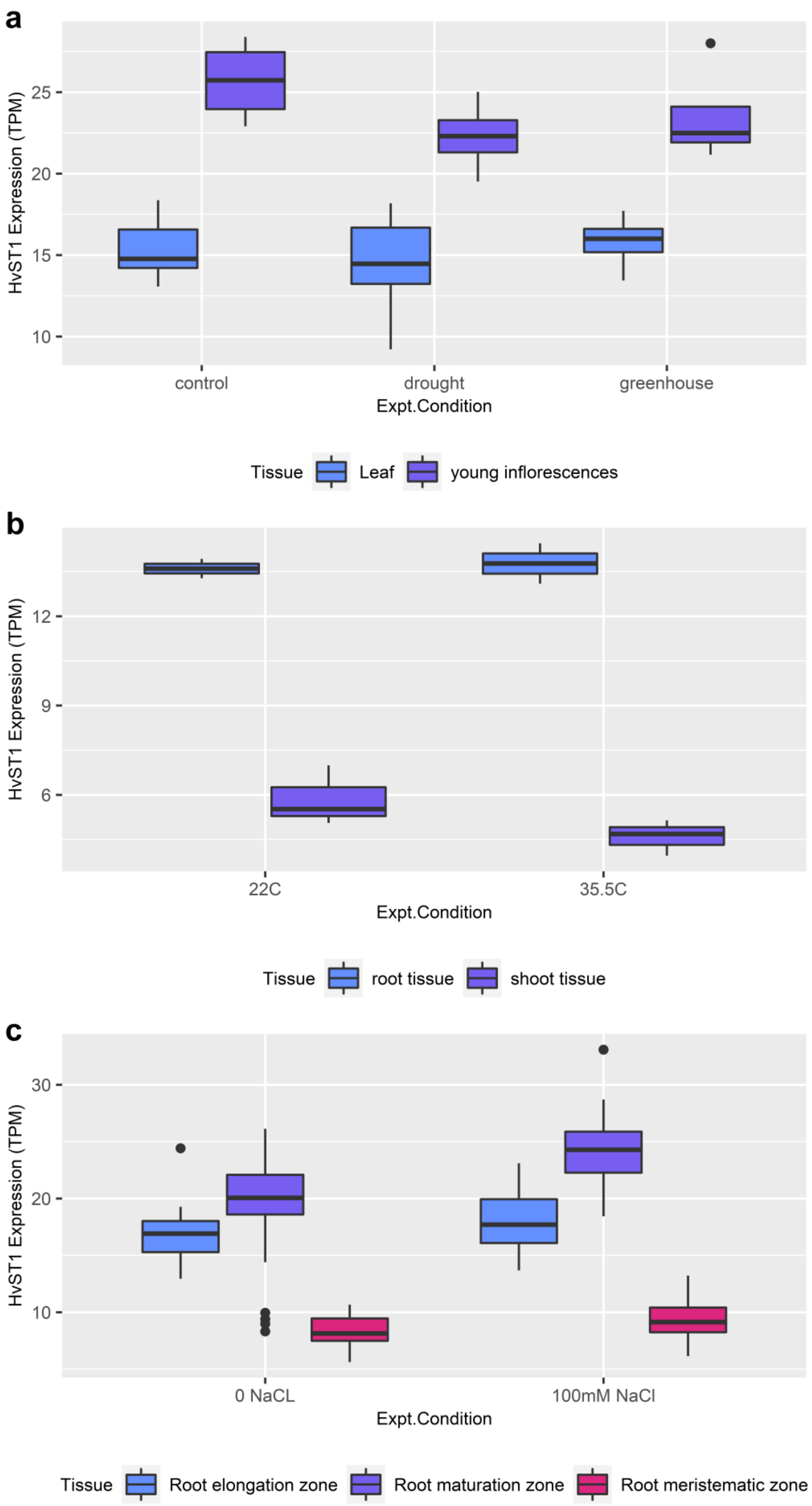


**Figure S10.** Boxplots of *HvST1* expression from the EoRNA database under drought (a), heat (b), and salt stress (c) conditions.

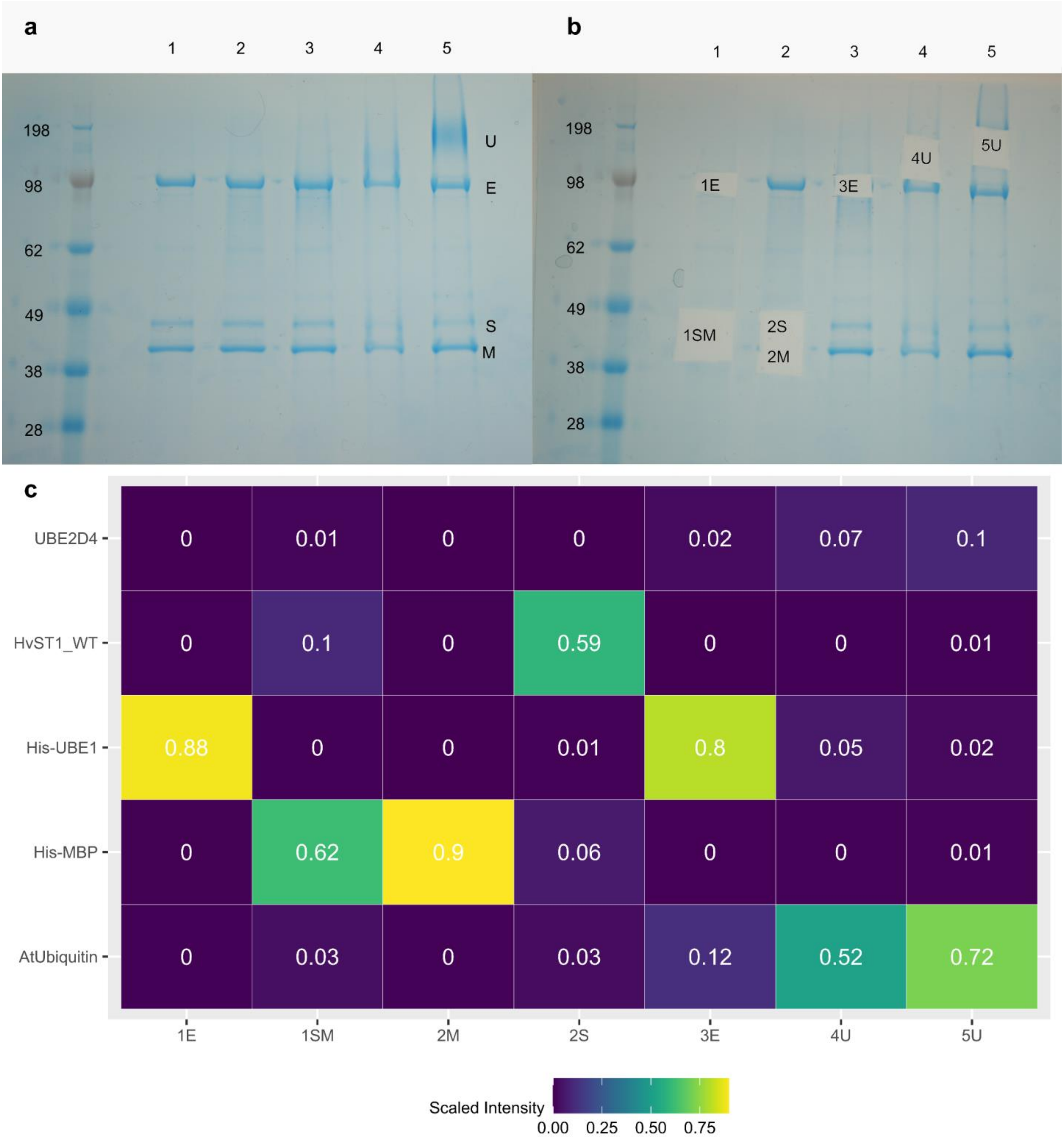


**Figure S11. Coomassie band identities.**
a) Coomassie stained SDS PAGE showing gel before and b) after dissection of fragments for in gel digestion and mass spectrometry to confirm the presence of Histidine tagged UBE1 (1E, 3E), Histidine tagged MBP (1SM, 2M), HvST1 (2S), and polyubiquitinated proteins (4U, 5U). c) The heat map shows the proportion of total protein intensity in mass spectrometry results for each dissected sample (x-axis) which confirmed the protein components of the autoubiquitination reaction (y axis).

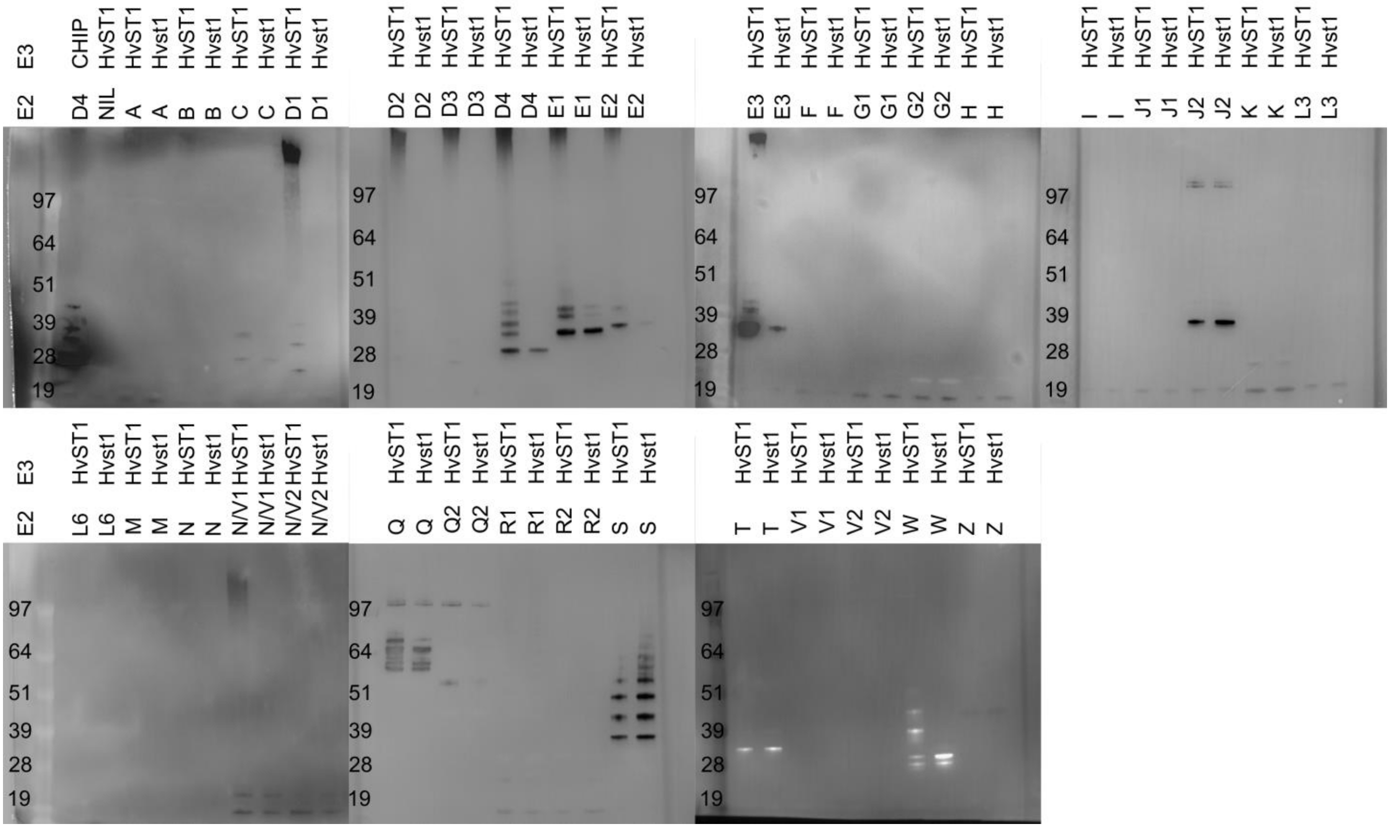


**Figure S12. E2 scan plate HvST1 vs *Hvst1***

Anti-ubiquitin conjugate antibody western blots showing HvST1 autoubiquitination in the presence of UBE2D and UBE2E family E2 conjugating enzymes and weaker interaction of HvST1 with UBE2A, UBE2B, and UBE2N/V1. Hvst1 did not interact with any E2 tested.

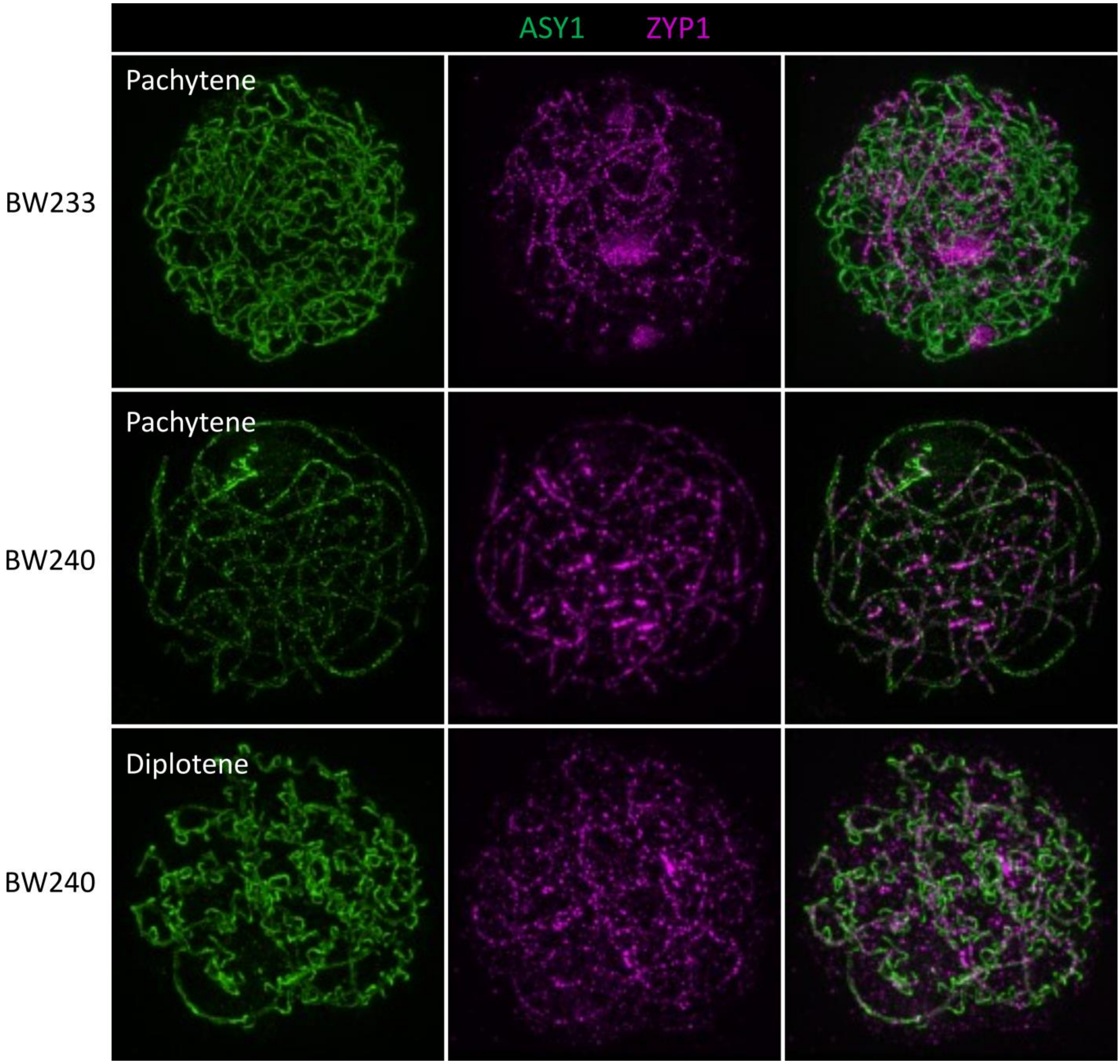


**Figure S13. late Synapsis of spontaneous *HvST1* mutants**

Pachytene and diplotene like cells of BW233 and BW240 which carries the same mutation in *HvST1.* ASY1 (green) and ZYP1 (magenta). Scale bar 5µm

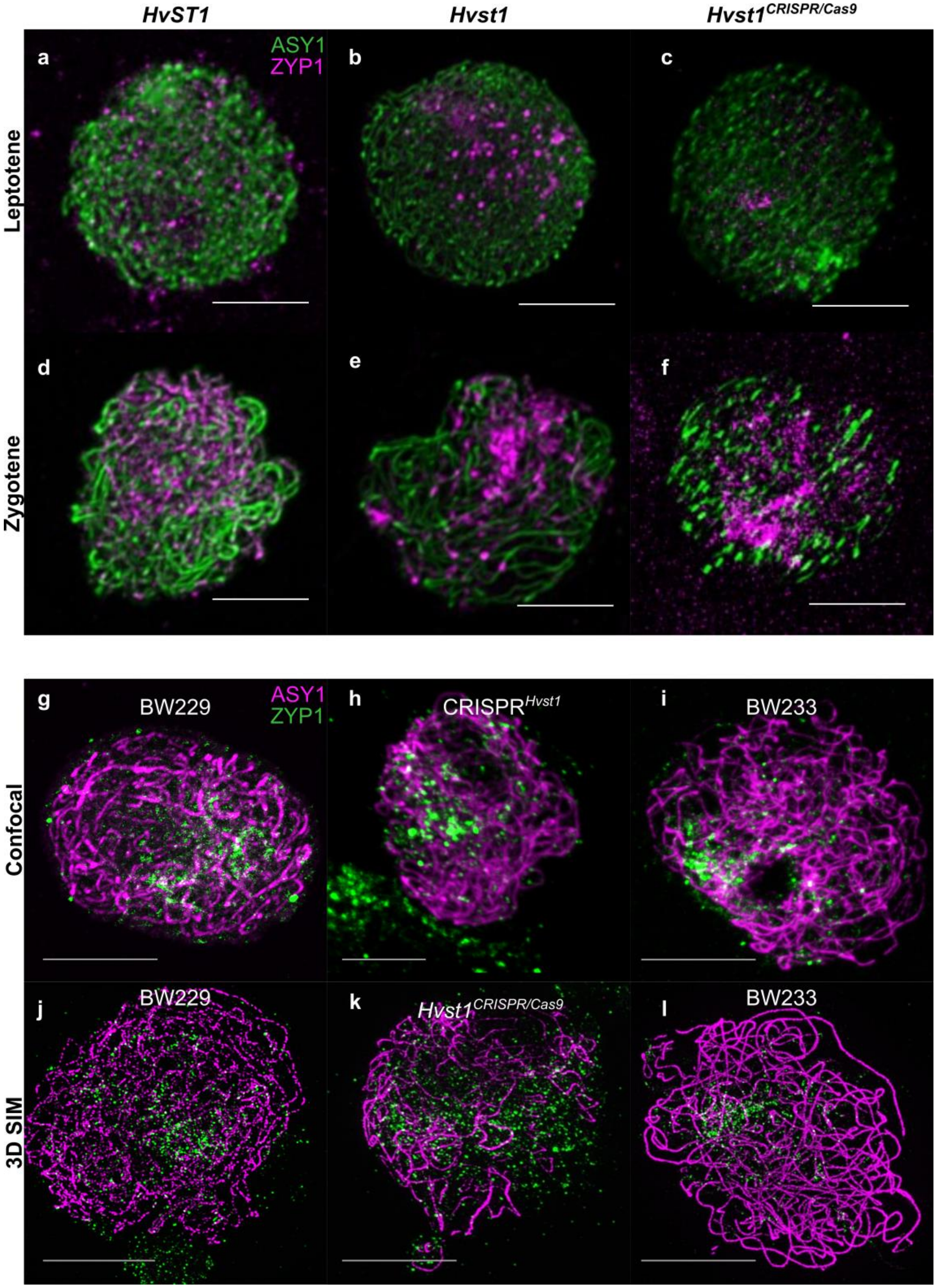


**Figure S14. Synapsis of spontaneous and induced *HvST1* mutants**

A- ASY1 and ZYP1, labelled in green and magenta respectively, were used to follow synapsis in a) d) Bowman (*HvST1*), b) e) BW233 (*Hvst1*) and c) f) CRISPR/Cas9 Line CLB2C18 (*Hvst1*$^{CRISPR/Cas9}$), which was designed to produce an *Hvst1 knockout*. a) At leptotene, cells of *HvST1*, b) *Hvst1,* and c) *Hvst1*$^{CRISPR/Cas9}$ look similar. d) At zygotene, synapsis progresses normally in *HvST1*, but ZYP1 forms a polycomplex in e) the spontaneous mutant *Hvst1* and f) the CRIPSR mutant *Hvst1*$^{CRISPR/Cas9}$ (*f*).

B- desynaptic mutants g,j) BW229 and l) BW240 exhibit the same abnormal synapsis phenotype as I,l) BW233. ASY1 (magenta) and ZYP1 (green). Scale bar 5µm

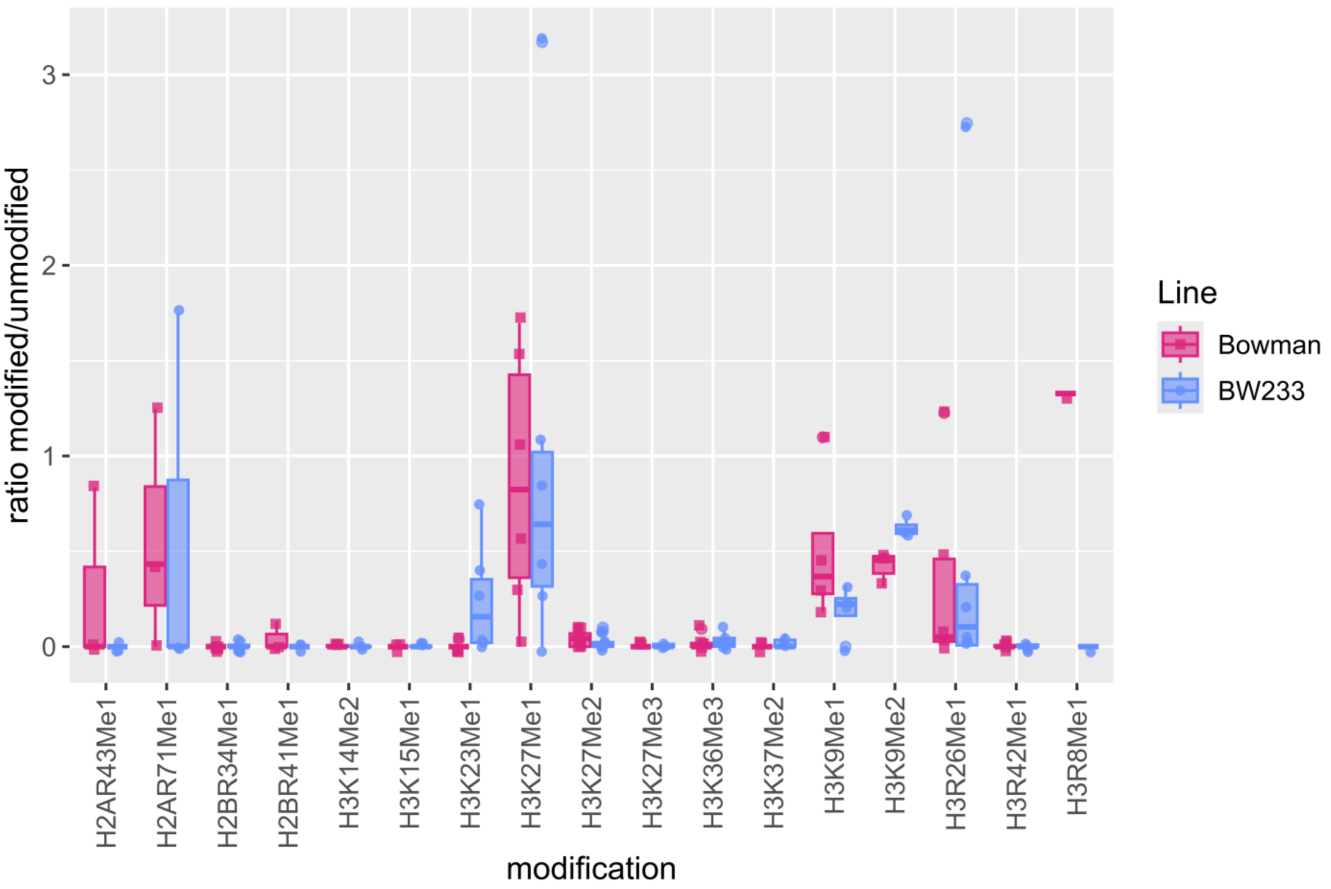


**Figure S15: Boxplot of histone methylation modification in WT and BW233**

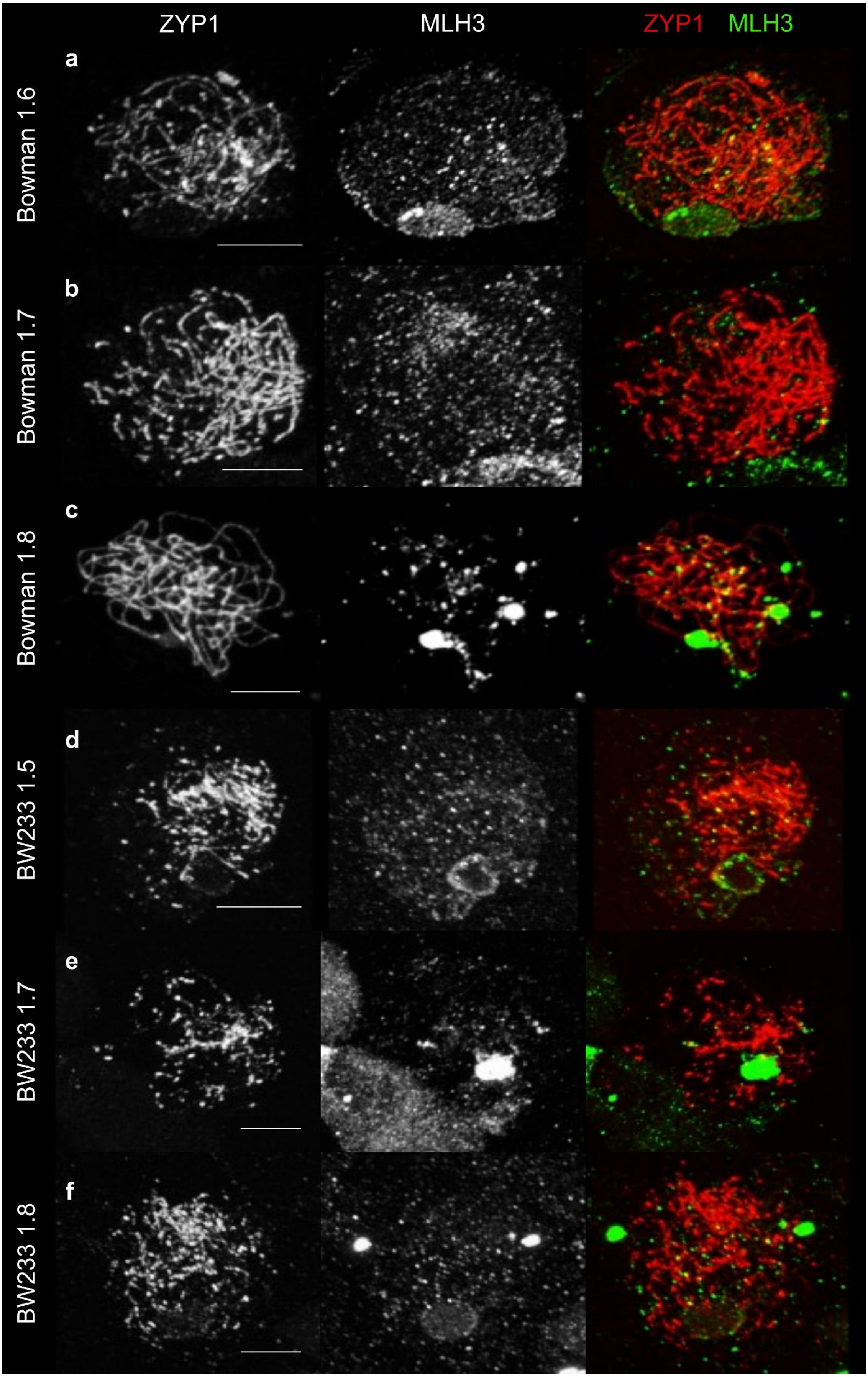


**Figure S16. MLH3 behaviour in Bowman and BW233**
a) b) In Bowman(*HvST1),* a large number of MLH3 (green) foci are present in the cytoplasm and on ZYP1 (red) axes during zygotene stage as previously described (Colas et al, 2016). c) At pachytene, large MLH3 foci locate the crossover sites. d) e) f) In BW233 (*Hvst1*), for a similar spike size, we see numerous MLH3 foci in the cytoplasm and the axes, but as we rarely reach a true pachytene, we cannot see many large MLH3 foci. Scale bar 5µm

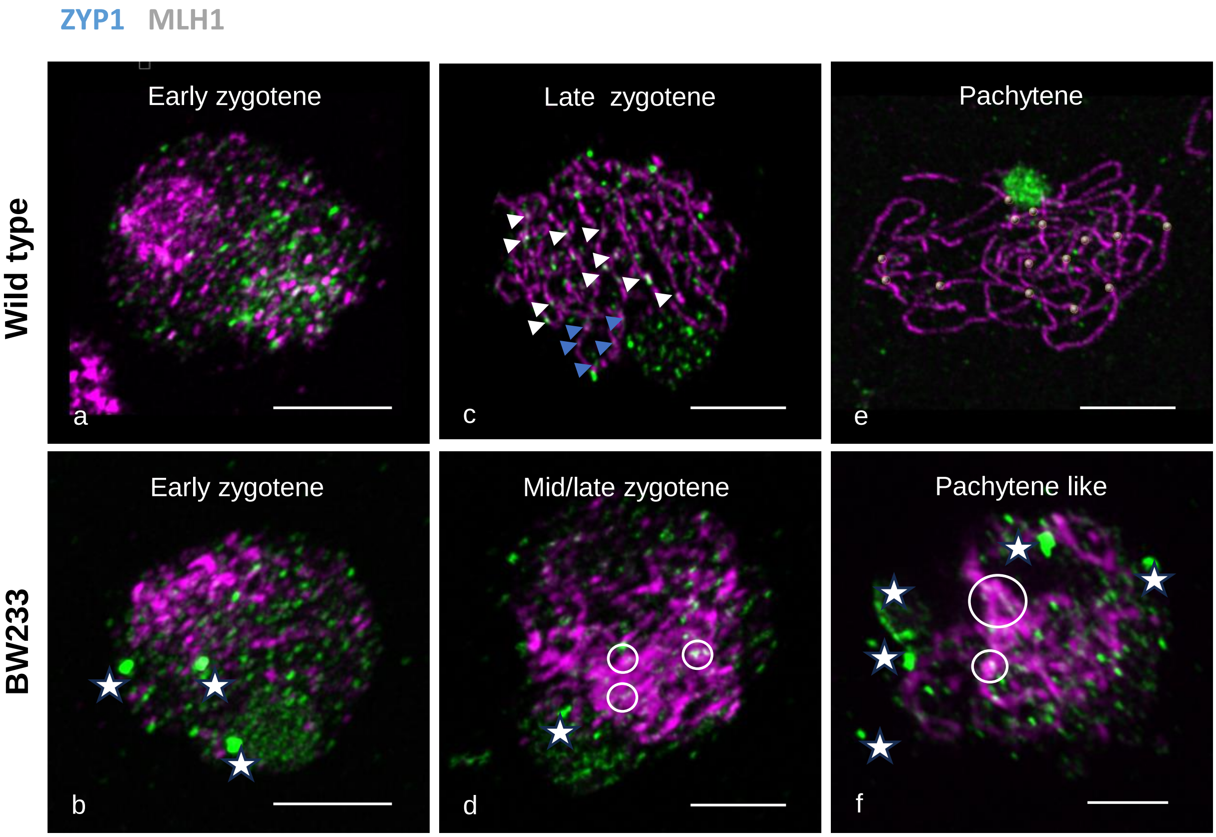


**Figure S17: Behaviour of MLH1 during synapsis**
Comparison of the Immuno co-localisation of ZYP1 (magenta) and MLH1 (Green) at a,b) early synapsis, c,d) mid synapsis and e,f) full synapsis of wild type and BW233. At the start of synapsis a), MLH1 relocalize from the cytoplasm towards the newly polymerized ZYP1 axes in the wild type. The same behaviour is observed in BW233 b), but we also see a lot more MHL1 polycomplexes (white stars). During synapsis elongation, MLH1 foci become more apparent on ZYP1 axes in the wild type c) presumably as a result of crossover formation (white arrows), but a lot of smaller foci (or intermediate) are also present (example with blue arrows). At pachytene, it is possible to count individual MLH1 foci in the wild type e). A lot of intermediates are also observed in BW233 d), but most of the MLH1 signal converges to the ZYP1 cluster and forms stretches rather than distinct foci (white circle). These stretches tend to be larger at a later stage f), similary to what we observed with our MLH3 antibodies. ZYP1 (magenta), MLH1 (Green), scale bar 5µm.

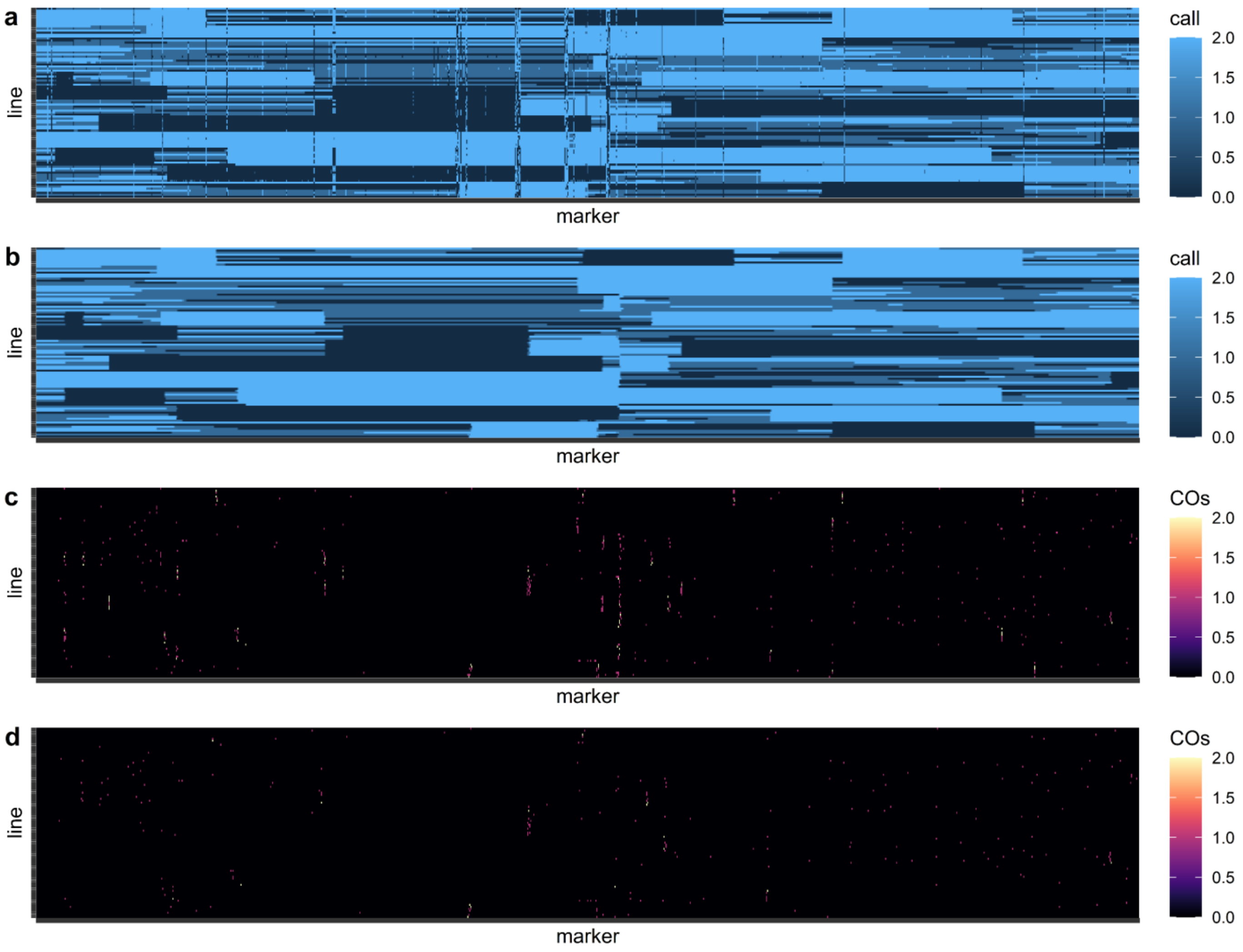


**Figure S18. Recombination Data Filtering**
(a) Heatmap of chr1H 50K iSelect markers in *Hvst1* lines (n=95) in order of physical position before and b) after sliding window correction indicating allelic state where 0 = match to Barke, 2 = match to Bw233, and 1 = heterozygous. c) Heatmap of chr1H crossover counts before and d) after filtering for F2 recombination events visible as large monomorphic blocks in panels a and b.
